# Selective collateralization of transcriptomically distinct neurons organizes visual-stream output from the primary visual cortex

**DOI:** 10.64898/2026.09.01.748541

**Authors:** Maryam Majeed, Mara C.P. Rue, Aixin Zhang, Sabrina Cheng, Jonathan Wang, Shannon Khem, Angela Ayala, Jeanelle Ariza, Olivia Helback, Katrina Nguyen, Ben Ouellette, Alana Oyama, Melissa Reding, Dean Rette, Julie Weber, Ali Williford, Yoh Isogai, Xiaoyin Chen

## Abstract

Parallel visual streams segregate information into pathways specialized for distinct computations, but how this segregation is achieved by anatomical segregation of primary visual cortex output remains unclear. This problem is complicated because individual neurons frequently send broadcasting projections to multiple cortical areas, and the target choice depends on both topographical location and molecular identity. Here we developed axonal BARseq2 to jointly map gene expression and high-resolution axonal projections from 1,448 neurons spanning the mouse primary visual cortex (VISp). Axonal BARseq2 recapitulated projection patterns observed by bulk tracing and single-neuron reconstruction, and recovered transcriptomic identities consistent with reference snRNA-seq datasets. Retinotopy strongly predicted projections to individual cortical targets, particularly for areas proximal to VISp, but explained little of which areas are frequently co-innervated. Instead, co-innervation patterns defined three preferential output pathways that largely corresponded to the ventral stream and two subdivisions of the dorsal stream. These pathways were associated with fine-grained transcriptional identities of L4/5 intra-telencephalic neurons, which were further validated with an external MERFISH dataset. Thus, VISp output is organized by two distinct rules: retinotopy constrains where neurons project, whereas cell-type-associated collateralization constrains which targets are co-innervated. This selective broadcasting, in which single neurons reach many higher visual areas in cell-type-specific combinations, could provide an anatomical substrate for visual-stream segregation at the level of VISp output in mice.

## Introduction

Visual information is processed through parallel cortical streams that emphasize distinct features of the visual scene and support different computations, including object recognition, motion and spatial processing, and visually guided action (Goodale and Milner, 1992; Nassi and Callaway, 2009). In primates, these functional streams are supported by pronounced anatomical specialization from the earliest stages of cortical processing: distinct pathways are segregated by cortical layer and module within the primary visual cortex (V1), and specialized populations in V1 and V2 preferentially route information toward downstream dorsal- and ventral-stream areas (Federer et al., 2009; Nassi and Callaway, 2009; Sincich and Horton, 2003). In the mouse, higher visual areas likewise exhibit distinct functional specializations and have been proposed to form analogous dorsal and ventral streams, with further subdivisions within the dorsal stream (Andermann et al., 2011; D’Souza et al., 2022; Marshel et al., 2011; Murakami et al., 2017; Wang et al., 2011, 2012; S. Yao et al., 2023). However, the anatomical segregation of these pathways at the level of V1 output is much less apparent. The mouse visual cortical network is densely interconnected (Harris et al., 2019; Wang et al., 2012, 2011). Projection-defined neuronal populations can be spatially intermingled within the primary visual cortex (VISp) (Kim et al., 2018), and individual VISp neurons commonly collateralize to multiple cortical targets(Han et al., 2018; Sorensen et al., 2026). Thus, although projection specialization into anatomical streams is evident across mouse higher visual areas, whether this segregation exists at the level of VISp output remains unclear.

Several organizing principles could underlie the segregation of visual streams at the level of VISp output. One possibility is topograph**y**. VISp and higher visual areas are organized into retinotopic maps, and projections from VISp to higher visual areas preserve systematic relationships between cortical location and visual space (Garrett et al., 2014; Marshel et al., 2011; Wang and Burkhalter, 2007; Zhuang et al., 2017). Consistent with a major role for spatial organization, recent single-cell reconstructions showed that a neuron’s tangential location within VISp strongly predicts its probability of projecting to particular cortical targets (Sorensen et al., 2026). Stream-specific projections could therefore arise, at least in part, from the superposition of spatially biased projections across the retinotopic map. Alternatively, stream organization could be reflected in which combinations of targets individual neurons innervate. Single-neuron tracing has shown that VISp neurons frequently send broadcasting projections to multiple cortical areas, but that these collateralization patterns are non-random (Han et al., 2018), raising the possibility that selective combinations of projections provide an additional level of organization beyond the probability of projecting to any individual target. Neuronal identity provides a further potential basis for segregating these outputs: broad cortical cell classes strongly constrain long-range connectivity (Harris and Shepherd, 2015), and finer molecular variation is associated with differences in projection patterns within the visual cortex (Sorensen et al., 2026). Determining how visual streams emerge from VISp output therefore requires systematically analyzing to what extent retinotopic topography and higher-order organization of projection collateralization distinctly contribute to individual projection probabilities, and asking whether the resulting output pathways are separable among molecularly defined neuronal populations.

Resolving these features simultaneously with existing approaches is challenging. Measuring the topography of long-range projections requires dense sampling and sufficient spatial resolution to link the locations of neuronal somata to their projection targets, whereas identifying cell types requires linking long-range projections to gene expression, all at single-cell resolution. To address this challenge, here we develop axonal BARseq2, which combines axonal BARseq and BARseq2, two in situ sequencing-based barcoded connectomics approaches, to jointly map the spatial, molecular and projection identities of individual VISp neurons. Axonal BARseq2 is based on BARseq (Chen et al., 2019), in which neurons are labelled with unique RNA barcodes that are amplified in the soma and transported along axons to axonal terminals, allowing the projection patterns of many individual neurons to be read out in parallel by sequencing. In axonal BARseq (Yuan et al., 2024), soma and axonal barcodes are simultaneously read out by in situ sequencing, enabling high-resolution mapping between soma and axon locations. BARseq2 (Sun et al., 2021) additionally captures the expression of endogenous genes together with neuronal barcodes, linking gene expression to projections in the same cells. The combination of these approaches therefore enables dense, single-cell measurement of projection topography, collateralization, and molecular identity within the same tissue, providing a direct means to determine how these factors jointly organize VISp output.

We applied axonal BARseq2 to map the projections and gene expression of 1,448 VISp neurons in two mice at single-cell resolution. Gene-expression and projection measurements agreed with previous single-nucleus RNA-sequencing (Z. Yao et al., 2023), bulk tracing (Harris et al., 2019; Oh et al., 2014), single-cell tracing (Sorensen et al., 2026), and functional retinotopic datasets (Waters et al., 2019; Zhuang et al., 2017). By resolving projections from individual neurons together with their spatial and transcriptomic identities, we found that projection targets depended strongly on tangential position within VISp, reflecting retinotopic biases that were most pronounced for nearby cortical targets. However, retinotopy did not explain which target areas were frequently co-innervated by the same neurons. Instead, collateralization patterns revealed three major output pathways, broadly corresponding to the ventral stream and two subdivisions of the dorsal stream, that were differentially associated with fine-grained L4/5 IT cell types. Thus, mapping single-neuron projections in their spatial and transcriptomic context reveals a cell-type basis for visual stream segregation in VISp.

## Results

### Axonal BARseq2 reveals diverse projections of mouse VISp neurons

BARseq-based techniques use padlock probe-based detection and amplification (Chen et al., 2018; Ke et al., 2013) combined with Illumina Sequencing-by-Synthesis chemistry (Chen et al., 2019, 2018; Sun et al., 2021) to capture both barcodes and endogenous genes at cellular resolution (**Fig. 1a**). Briefly, target mRNAs are reverse transcribed and hybridized with padlock probes. For endogenous genes, the two arms of padlock probes bind precisely to each other and are ligated, followed by rolling circle amplification. For random barcodes, the two arms bind to the constant regions that flank the barcode, and a DNA polymerase is used to copy the barcode into the padlock probe (i.e., “gap-filling”). The gap-filled padlock probe is then ligated and amplified as for endogenous mRNAs. There are two technical challenges in combining in situ sequencing of endogenous genes (i.e., BARseq2) (Sun et al., 2021) and axonal barcodes (i.e., axonal BARseq) (Yuan et al., 2024). First, the gap-filling polymerase reduces the detection of endogenous genes, possibly because of its proofreading activity. In BARseq2, the polymerase is highly diluted to achieve a balance between gene and barcode detection, but is less robust for either. To boost detection of both endogenous genes and barcodes, we performed two rounds of rolling circle amplification using two phi29 DNA polymerases with complementary characteristics: an engineered phi29-XT DNA polymerase that can generate more, but weaker amplicons, and the wild type phi29 DNA polymerase that robustly produce rolling circle amplicons with bright signals. Together, tandem application of these two phi29 DNA polymerases allows the original phi29 to continue amplifying the weak amplicons achieved by phi29-XT, thereby achieving higher counts of endogenous genes per cell compared to the original BARseq2 protocol (**ED Fig. 1a-c**). Second, capturing axonal barcodes requires sequencing many sections to cover a substantial portion of the mouse brain, which is both time consuming and error-prone. This challenge is already present in axonal BARseq (Yuan et al., 2024), but sequencing both endogenous genes in addition to axonal barcodes results in 1.5× the number of sequencing rounds and further amplifies the challenge. To make sequencing faster and more robust, we used a custom in situ sequencing platform to fully automate the sequencing reactions and imaging. Together, these modifications allowed axonal BARseq2 to robustly sequence both endogenous genes and barcodes in both somata and axons across a substantial portion of the mouse brain.

**Figure 1.**
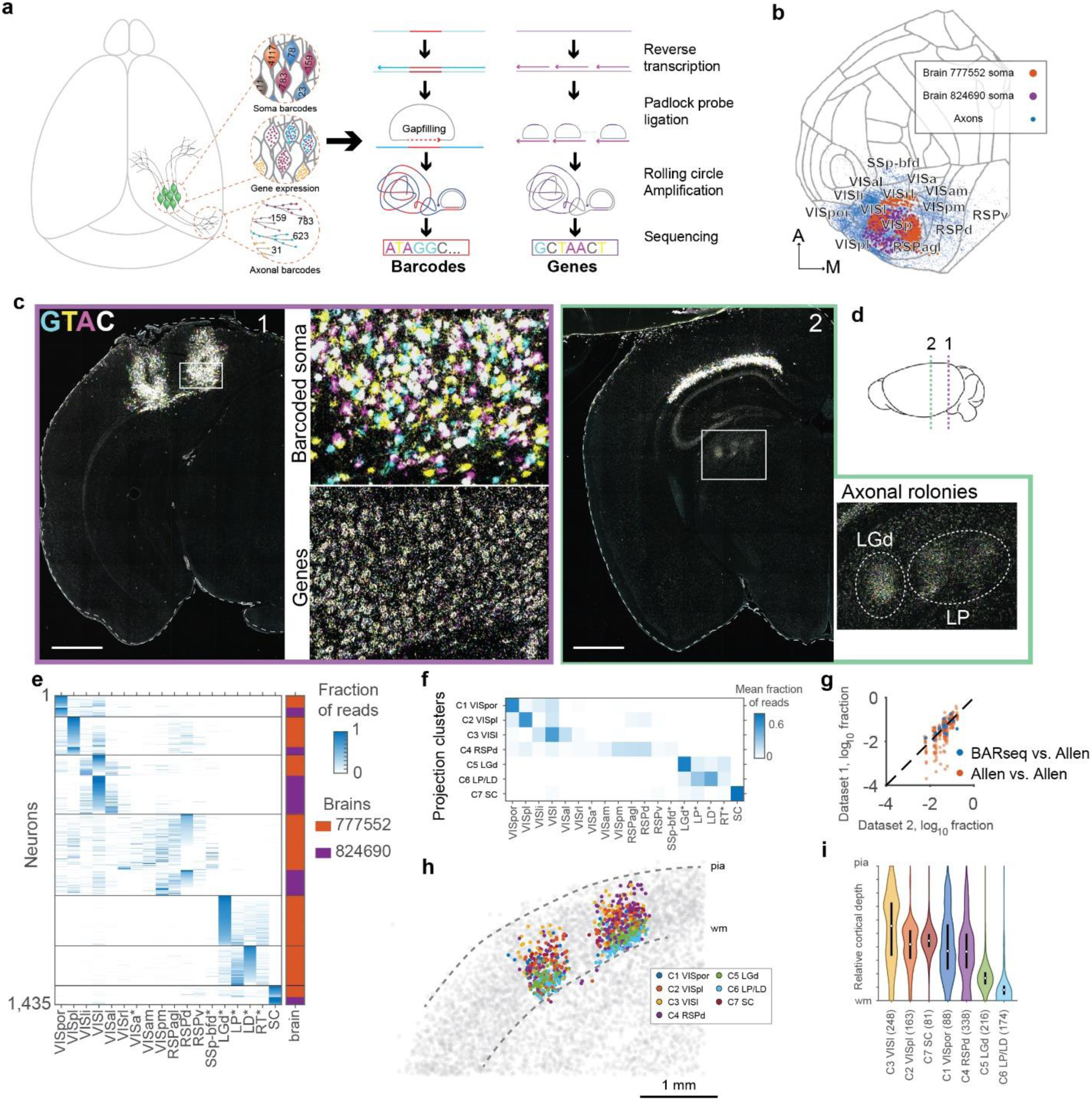
Axonal BARseq2 reveals diverse VISp projections. (a) Schematic illustrating axonal BARseq2. Barcodes and endogenous genes are sequenced in somata and axons; both barcodes and genes are amplified and sequenced in situ using a padlock probe-based approach. (b) Barcoded somata and axonal barcodes visualized on a cortical flatmap. Big and small dots indicate soma and axonal barcodes, respectively; color indicates brains. (**c**)(**d**) Representative images from the first barcode sequencing cycle on two hemi-coronal sections. Zoomed-in views show somata barcodes and endogenous genes in the VISp region, and axonal barcodes in the thalamus. Two thalamic nuclei (LGd, LP) with dense axonal barcodes are shown. The nucleotides that correspond to the colors are indicated. Scale bars = 1 mm. Locations of the two hemi-coronal sections are shown in (d). (**e**) Heatmap showing the row-normalized projections of individual neurons (rows). Columns indicate projection strengths to the indicated target areas. Color bar on the right indicates brains. * indicates areas that were only sequenced substantially in brain 777552. (**f**) Heatmap showing the mean projection strengths of neurons in each projection cluster (rows) to each target area (columns). (**g**) Comparison of area-to-area connectivity between this dataset (dataset 1) and the Allen Mouse Brain Connectivity Atlas (dataset 2, blue), and between different experiments in the Allen Mouse Brain Connectivity Atlas (orange). (**h**) Barcoded somata colored by their projection clusters from 20 stacked sections. Gray dots indicate non-barcoded somata. (**i**)Violin plot showing the distribution of laminar positions of neurons from distinct projection clusters.

We then applied axonal BARseq2 to map the transcriptomic identity and projections of neurons from the primary visual cortex (VISp) of two mouse brains. In each brain, we injected barcoded Sindbis virus at five and six sites, respectively, that tiled VISp (**Fig. 1b**); across the two brains, the labeled neurons provided complementary spatial coverage, together spanning nearly the entire VISp. We sectioned the brains to 20µm hemi-coronal sections, and every other slice was processed for in situ sequencing (**Fig. 1c, d**) to read out both barcodes and expression of 104 cortical excitatory neuron marker genes (Chen et al., 2025). These sections spanned the posterior 2.6 mm and 1.9 mm of a cortical hemisphere in each brain, respectively, totaling 112 hemi-sections across the two brains. The sampled projection areas covered major cortical projection targets of VISp, including VISam, VISpm, VISrl, VISal, VISl, VISli, VISpor, and RSP (**Fig. 1b**) and the superior colliculus (SC); one of the brains additionally included anterior cortical regions SSp-bfd, VISa, and thalamic nuclei including the dorsal lateral geniculate nucleus (LGd), the lateral posterior nucleus (LP), and the posterior half of the lateral dorsal nucleus (LD). Altogether, these areas covered most major projection targets of VISp, except for the striatum (Harris et al., 2019). We processed the gene expression data following our previously established pipeline (Chen et al., 2025). All data were registered to the Allen Common Coordinate Framework (CCF) as previously described (Chen et al., 2025) using QuickNii and Visualign (Puchades et al., 2019).

Because Sindbis-encoded barcodes are highly expressed, barcoded somata are often filled with barcode mRNA rolonies that blur together into one blob, whereas barcode molecules in the axons show up as clouds of individual rolonies (**Fig. 1c**). We detected barcoded somata based on barcode signal intensity and manually proofread the cells to exclude barcodes in neighboring cells and/or overlapping axons and dendrites to be assigned to the wrong cells. Barcodes in somata and rolonies in the axons were decoded by identifying the channel with the strongest signal, as previously described (Zhang et al., 2024). We then matched the axonal barcodes to soma barcodes, allowing at most one mismatch (see **Supp Note 1** and **ED Fig. 2a-c** for validation of barcode matching). We focused on 1,448 neurons whose somata were localized to VISp, and had sufficient barcodes detected in the axons (≥5 rolonies outside of VISp) for downstream analysis (**Fig. 1d**). None of the 1,448 soma barcodes were shared across the two brains, confirming that the barcode diversity was sufficient for single-cell resolution.

**Figure 2.**
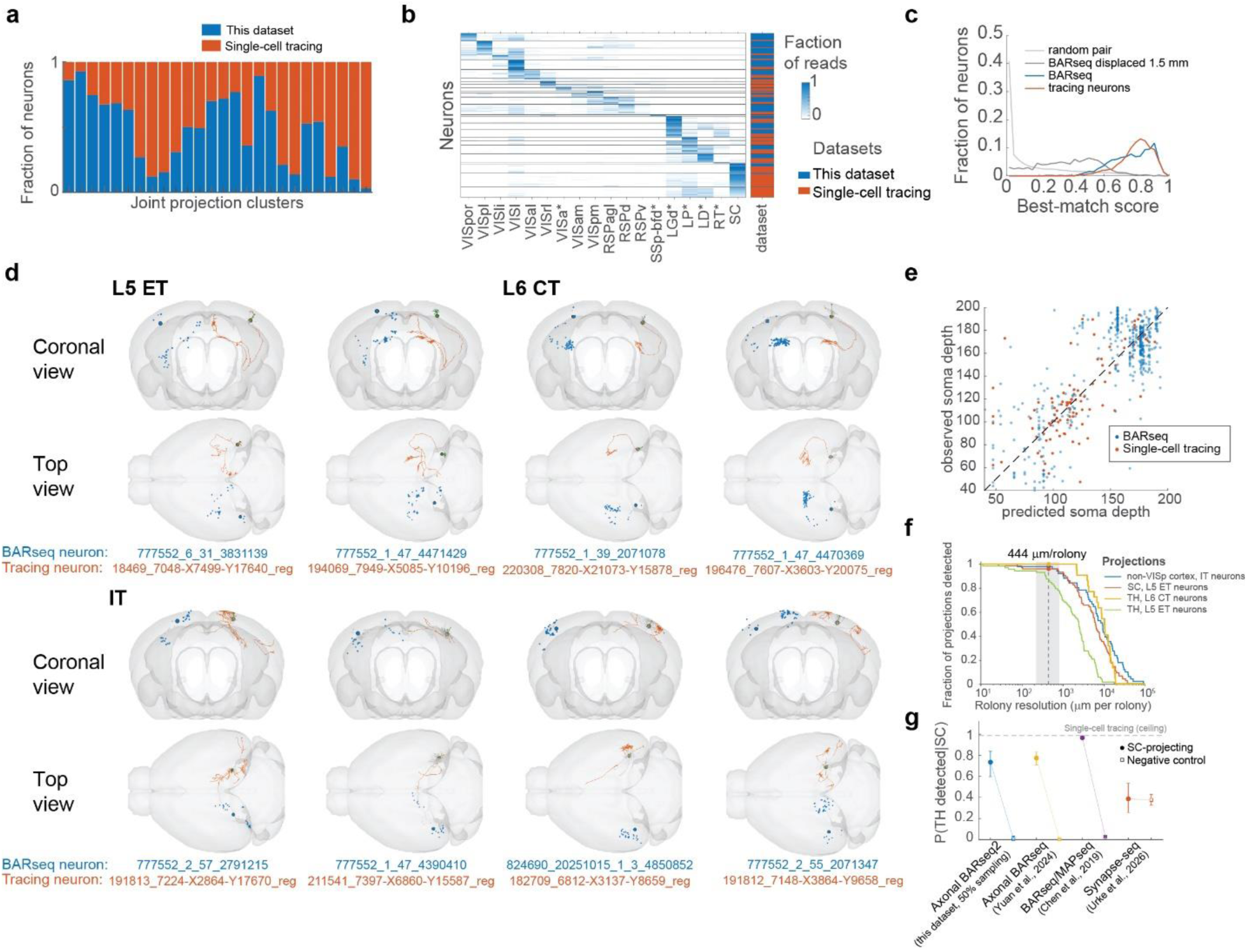
Axonal BARseq2 recapitulates single neuron tracing data. (**a**)(**b**) Hierarchical clustering of combined traced neurons (Sorensen et al., 2026) and this dataset after matching the sizes of the two datasets, resulting in 26 clusters each containing neurons from both datasets. (a) shows the fraction of neurons in each joint cluster, and (b) shows the row-normalized projection matrix, sorted by cluster identity. Clusters are separated by horizontal lines in the projection matrix. (**c**) The distribution of MCS for best matches to neurons in this dataset, for best matches to neurons in this dataset after displacing them by 1.5mm, for best matches to traced neurons, and MCS for random pairs of a neuron in this dataset and a traced neuron. (**d**) Example neurons from this dataset (blue) plotted with the best matched traced neuron (red). In each brain, the traced neuron has been flipped to the right hemisphere and plotted only projections on the ipsilateral side. The neuron IDs are listed at the bottom of each pair of neurons. For each traced neuron, the dendrites are plotted in green. For BARseq neurons, somata are plotted with a big blue dot, and axonal barcodes are shown as small blue dots. (**e**) The actual soma depth (y axis) of a BARseq neuron (blue) or a traced neuron (red) compared to its predicted soma depth (x axis) based on the mean depth of its top three matches in the traced neuron dataset. Dashed line indicates diagonal. (**f**) Fraction of axons detected (y axis) at the indicated rolony sensitivity (x axis, in µm of axons per rolony). The dashed vertical line indicates the estimated sensitivity in this experiment, and the gray area spans the estimated sensitivity of the technique (220 µm) and the detection floor observed in the TH collaterals of SC-projecting neurons (810 µm). (**g**) Fraction of TH collaterals in SC-projecting neurons detected in each dataset (solid dots) and in negative control neurons that should not have TH projections (squares). Error bars indicate Wilson 95% confidence interval for this dataset, Yuan et al. 2024, and Chen et al. 2019, and 95% interval from Monte Carlo propagation of uncertainty through the collision correction in Urke et al., 2026. The negative control neurons are superficial layer, non-SC projecting neurons in this dataset, no-SC projecting, contralaterally projecting neurons in Yuan et al., 2024 and Chen et al., 2019, and transcriptomically defined glial cells in Urke et al., 2026.

VISp neurons projected predominantly to classical higher visual areas, retrosplenial areas (RSPd, RSPv, and RSPagl), and the primary somatosensory cortex (SSp); subcortically, neurons projected to the thalamic nuclei LGd, LP, and LD, and the superior colliculus (SC) (**Fig. 1e**). To examine whether the projections in our dataset were consistent with bulk anterograde tracing results, we computed pseudo-bulk projection strengths by aggregating BARseq projections across neurons for each target area (**Fig. 1f**) and compared them to projection patterns described in the Allen Connectivity Atlas (Harris et al., 2019; Oh et al., 2014). Our data showed strong correlation with bulk tracing data (**Fig. 1g**; Pearson r = 0.68, p = 0.002 by permutation); this correlation was comparable to the variability observed across different injection sites in VISp in the Allen Connectivity Atlas itself (Pearson r = 0.71, p = 0.0001 by permutation). These results demonstrate that our data recapitulated bulk-level projection patterns of mouse VISp neurons.

### Axonal BARseq2 recapitulates projection patterns from single-cell reconstruction

To analyze projections at single-cell resolution, we first performed a coarse-level hierarchical clustering to reveal broad patterns of projections. We found that two clusters of neurons in L6 (C5 LGd, C6 LP/LD) projected to LGd and LP/LD, respectively; one cluster (C7 SC) of L5 neurons projected to the SC; and the remaining clusters (C1-C4) of neurons across cortical layers projected to various combinations of cortical areas (**Fig. 1e, h, i**; **ED Fig. 2d** for clustering stability). These broad patterns were consistent with the classic divisions of IT, PT, and CT neurons (Harris and Shepherd, 2015). For areas sampled in both brains, neurons showed similar projection patterns (**Fig. 1e**; **ED Fig. 2e, f**). To assess whether our data captured projection patterns of single neurons at high resolution, we compared our data to 169 published single neurons in VISp that were traced individually (Sorensen et al., 2026). To compare the two datasets at the resolution of target areas, we resampled the traced neurons to match the higher number of neurons and the lower axonal resolution of the BARseq dataset, and combined 1,435 such “BARseq-like neurons” with 1,435 real BARseq neurons that had projections in the 18 areas we compared. We then over-clustered the combined data and found that no cluster was exclusive to either dataset, and this held true up to 26 clusters.(**Fig. 2a,b**). Despite enrichment of different subpopulations of neurons across the two datasets, these results suggest that individual neurons were similar across the two datasets at the resolution of target areas. To assess consistency beyond area resolution, we matched individual BARseq neurons to the traced neurons. We defined a “Match Consistency Score” (MCS) between a query single neuron (either a BARseq neuron or a traced neuron) and a reference neuron (traced neurons from Sorensen et al., excluding the query neuron), as the harmonic mean of optimal transport that transforms the axonal data between the two neurons (see **Methods** for details). MCS ranges from 0 (no matching axon) to 1 (perfect match). Because the soma locations and dendrite information are both excluded from MCS, MCS reflects the consistency of the spatial distribution of axons, rather than other anatomical features of the neuron. We then identified the best match reference neurons for each BARseq and traced neuron (**Fig. 2c, d**). The best matched traced neurons had a MCS of 0.80 ± 0.10 (mean ± s.d.), whereas BARseq neurons had a MCS of 0.77 ± 0.13 (mean ± s.d., 3.5% below the ceiling defined by traced neurons). As controls, we shifted each BARseq neuron by 1.5 mm and re-identified the best matched neurons in the tracing dataset. The best-matched neurons for the shifted BARseq neurons had an MCS of 0.38 ± 0.19 (mean ± s.d., p = 1×10^−179^ compared to the best matches to the unshifted BARseq neurons, paired signed-rank test) (**Fig. 2c**), demonstrating that MCS is sensitive to where axons are located and can distinguish real matches from spurious ones.

Visual inspection of the matched neurons revealed that our data captured diverse projection patterns both within and across broad classes of projections (L6 CT/L5 ET/IT) (**Fig. 2d**). For example, Some L5 ET neurons branch more extensively in the thalamus (e.g. 777552_1_47_4471429) compared to others (777552_6_31_3831139). Among L6 CT neurons, some neurons (e.g., 777552_1_39_2071078) projected more focally and laterally, whereas others (e.g., 777552_1_47_4470369) project more dorsally and medially, covering a broader region in the thalamus (**Fig. 2d, L6 CT examples**). Across the IT neurons, some neurons projected focally to neighboring areas (e.g., 824690_20251015_1_3_4850852), whereas others projected only to deep layers of the cortex (e.g., 777552_1_47_4390410). These detailed anatomical differences were consistent with those seen in the best matched traced neurons.

As an independent test of the correspondence between BARseq and traced neurons, we asked whether axonal similarity alone could predict soma depth, which was excluded from the calculation of MCS. Specifically, we predicted the soma depth of each BARseq neuron from its top three well-matched traced neurons (MCS > 0.8, **Fig. 2e**). The predicted soma locations of BARseq neurons were correlated with the real soma location (Pearson r = 0.75, p = 4×10^−89^) to a similar extent as the predicted and real soma locations of traced neurons (Pearson r = 0.70, p = 4×10^−16^), suggesting that the remaining uncertainty likely reflected biological heterogeneity and/or lack of coverage in the reference tracing dataset. These results indicate that the axonal BARseq2 data were highly consistent with single-cell tracing data.

To assess the sensitivity of axonal BARseq2 in mapping axons, we examined the distribution of branched axon length and barcode counts in the same projections. We used the thalamic (TH) collateral of the SC-projecting neurons as the reference projection, because it could be detected robustly in the single-cell tracing dataset (73 of 76 SC-projecting neurons; see Methods). This results in a conservative estimate of BARseq sensitivity, because the in situ sequencing only covered the posterior half of LD and is thus missing a substantial amount of thalamic axons due to the partial tissue coverage. The BARseq-detected TH collaterals of the SC-projecting neurons had a median of 6.5 rolonies, compared to a median of 2.9 mm of branched axon length in traced neurons, suggesting that in our dataset one rolony on average corresponds to 440 µm of axons. Because we sequenced every other section in this dataset (i.e., 50% sampling rate), the resolution limit of axonal BARseq2 itself is 220 µm of axons per rolony. Based on the distribution of axon lengths observed in the traced neuron dataset, we modeled detection of axons in traced neurons by rolonies given a range of sensitivity (from 220 µm to 814 µm) that straddles the estimated value. This range of sensitivity is sufficient to robustly detect major projections, including 92-98% of non-VISp cortical projections of IT neurons, 100-100 % of thalamic projections of L6 CT neurons, and 93-96% of SC projections of L5 ET neurons, and 78-95% of TH collaterals of L5 ET neurons (**Fig. 2f**; see Methods).

To compare the sensitivity of our technique to other barcoded connectomics techniques, we used the detection probability of TH projections in SC-projecting neurons as a reference metric (**Fig. 2g**). The TH collaterals of SC-projecting neurons are detected at a comparable rate in this dataset (mean 73%, 95% CI 60% – 84%) and the original axonal BARseq dataset (Yuan et al., 2024) (mean 77%, 95% CI 71-83%, p = 0.6 compared to this dataset using Fisher’s exact test) despite the 50% section sampling plan, whereas BARseq and/or MAPseq with bulk sequencing at the projection sites (Chen et al., 2019) detected the TH collaterals at a higher rate that is largely comparable to single-cell tracing (mean 97%, 95% CI 96-97%). All three approaches had negligible detection in negative control populations that do not project to the thalamus (0.6% for this dataset, 0.4% for the original axonal BARseq, and 3% for bulk sequencing at projection sites). All three Sindbis virus-based approaches performed markedly better than the AAV-based Synapse-seq (Urke et al., 2026), which detected TH collaterals in 39% of SC-projecting neurons, compared to a noise floor of 38% in glial cells. Altogether, these results indicate that our dataset can detect most projections of VISp neurons and achieve sub-area and laminar resolution, though it may miss a small fraction of weak projections.

### Single-neuron VISp projections recapitulate the retinotopy of VISp projections

Recent single-neuron reconstructions suggested that neuronal identity and cortical position jointly constrain projection targets in VISp (Sorensen et al., 2026). However, the extent to which this spatial dependence reflects retinotopy, and how strongly retinotopic position structures both target selection and the organization of projections within targets, remain unclear. To address these questions, we investigated the spatial relationship between VISp somata and their projections. Retinotopic maps vary substantially across individual animals (Waters et al., 2019). To estimate the retinotopic representation of neurons in our dataset, we calculated an average retinotopic map based on intrinsic signal imaging data across 283 animals (Harris et al., 2019; Oh et al., 2014; Waters et al., 2019) in CCF space, then assigned the altitude and azimuth values to each BARseq neuron and its associated axonal barcode rolonies based on its soma location (**ED Fig. 3a**).

**Figure 3.**
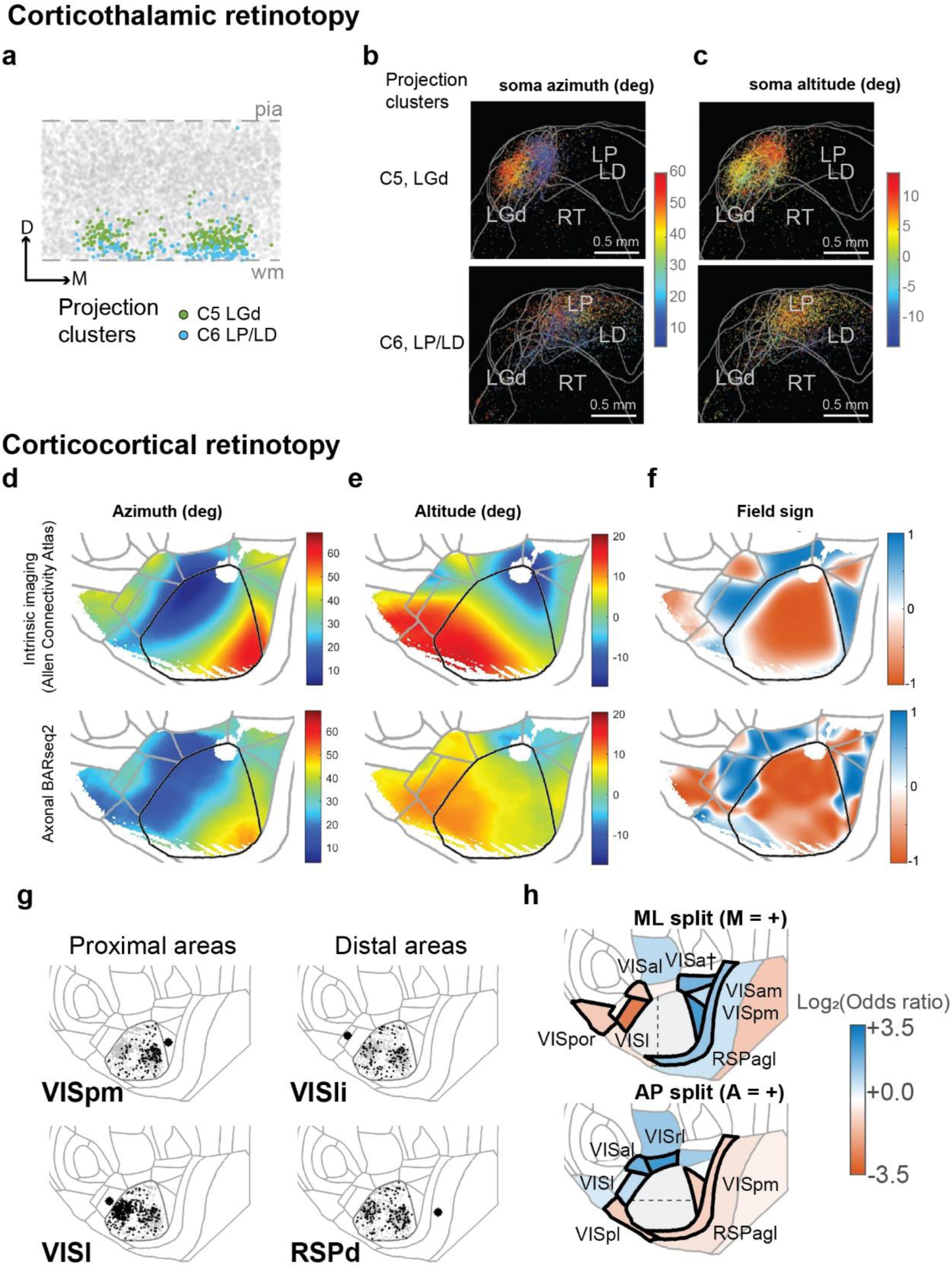
Axonal BARseq2 recapitulates visual cortex retinotopy. (**a**) The locations of C5 and C6 cluster neurons shown in flatmap “coronal” view. X axis indicates the ML axis of the flatmap, and y axis indicates the depth axis of the flatmap. (**b**)(**c**) Projection rolonies of C5 and C6 neurons, collapse onto a single coronal plane. The approximate locations of large thalamic nuclei are drawn. Axonal barcodes are colored by azimuth (b) and altitude (c) as indicated. (**d**)-(**f**)Azimuth (d), altitude (e), and field sign (f) maps from intrinsic imaging data in the Allen mouse brain connectivity atlas (top row) and from axonal barcodes of this dataset (bottom row). For this dataset, each axonal barcode inherits the azimuth and altitude values of its soma as inferred from **ED.** Fig. 3a. In all six plots, only locations where sufficient rolonies were present to accurate retinotopy were shown. (**g**) The locations of neuronal somata (black dots) that project to the indicated target cortical areas. Neurons that do not project to the target areas are shown in gray, and the relevant target areas are shown with an asterisk. The left column contains proximal areas, and the right column contains distal areas. (**h**) Spatial biases in projections to each target area along the mediolateral (top) and anteroposterior (bottom) axes of VISp. Color indicates the log₂ Mantel–Haenszel odds ratio for projection to each target from neurons with somas in the medial half (*top*) or anterior half (*bottom*) of VISp, stratified by brain and using the Haldane–Anscombe correction. Black outlines indicate FDR < 0.05, and the significant area names are labeled.

We first examined the retinotopic organization of corticothalamic projections. Of the two clusters of L6 CT neurons, C6 projected mainly to LP/LD with somata at the bottom of L6, and C5 projected mainly to LGd with somata at a more superficial position in L6 (**Fig. 3a**). Within each cluster, neurons projected to different locations in an orderly fashion that reflected the retinotopic organization of their somata (**Fig. 3b, c**). In the C5 LGd-projecting cluster, azimuth increased along the medial-to-lateral axis, whereas altitude increased along the ventral-to-dorsal axis. In the C6 LP/LD-projecting cluster, azimuth varied primarily along the ventrolateral-to-dorsomedial axis (**Fig. 3b, c**). Because altitude in LP is typically organized along the anterior-posterior axis (Allen et al., 2016), and the anterior extent of LP/LD was incompletely sampled in our dataset, we were unable to resolve a clear altitude gradient. These results indicate that corticothalamic projections from the two projection-defined subpopulations form distinct retinotopic maps in LGd and LP/LD, consistent with previous mapping studies (Allen et al., 2016; Bennett et al., 2019; Born et al., 2021; Piscopo et al., 2013).

We next examined the retinotopic organization of corticocortical projections of the IT neurons. For each location across the visual cortex, we calculated the weighted average of projection rolonies in that local area, then rescaled the projection-inferred retinotopy to match the original scale (**Fig. 3d, e**). The projection-inferred azimuth, altitude, and the field sign map showed patterns that resembled the functionally defined maps across the whole visual cortex (**Fig. 3d-f**). To assess the agreement between the projection-inferred and functional retinotopic maps, we calculated their pixel-wise correlation and compared it with correlations between individual brains and the average of the remaining brains in the Allen Connectivity Atlas dataset. The projection-inferred maps had a Pearson correlation of 0.778 for azimuth and 0.783 for altitude, both of which fell within the 95% confidence interval of the correlation across individual brains (95% CI: azimuth 0.62 to 0.91, p = 0.29; altitude 0.70 to 0.92, p = 0.23) (**ED Fig. 3b, c**). These results indicate that the projection-inferred retinotopic map is within the range of variation across individuals.

Because different higher visual areas represent different portions of visual space, this organization further predicts that retinotopic position within VISp should influence which cortical areas they target. Consistent with the recent finding that cortical position within VISp predicts projections to higher visual areas (Sorensen et al., 2026), we found that the spatial dependence of projection target preference largely reflected the portion of visual space represented by each higher visual area. Across higher visual areas, mean azimuth and altitude inferred from projections were strongly correlated with the corresponding means from the functional maps (**ED Fig. 3d, e**; Pearson r = 0.82, p = 0.006 for azimuth and r = 0.88, p = 0.002 for altitude). For example, VISl functionally represented more medial portions of visual space and received projections predominantly from the lateral half of VISp, whereas VISpm represented more lateral portions of visual space and received projections predominantly from the medial half of VISp (**Fig. 3g**; **ED Fig. 3f**). Strikingly, this dependence of target selection on soma position was especially prominent among cortical areas proximal to VISp. All proximal areas showed significant enrichment along either the mediolateral or anteroposterior axis (**Fig. 3h**, black outlines; FDR < 0.05), whereas among distal areas (including VISli, VISpor, RSPd, RSPv, and SSp-bfd), only VISpor showed a significant spatial bias. Other distal areas, such as RSPd and VISli, were comparatively insensitive to soma position (**Fig. 3g**). Thus, retinotopy not only organizes projections within cortical target areas, but is also a major determinant of how VISp output is routed among VISp-proximal cortical areas. This correspondence raises the possibility that target-specific projection patterns, including biases in collateralization across higher visual areas, could arise largely from the partial representation of visual space in each target area.

### VISp single neurons preferentially collateralize along three pathways

The classic theory of the mouse visual system separates higher visual areas into two groups that are reminiscent of the primate dorsal stream and ventral stream. Han et al. (Han et al., 2018) previously found that VISp neurons are more likely to project to specific combinations of higher visual areas, but whether such preferences in collateralization reflect the visual processing stream remains unclear. We first tested whether our data could recapitulate the same enriched collateralization patterns. Following Han et al. (Han et al., 2018), we modeled each projection as a binomial distribution, and identified combinations of projections that were over- or under-represented in the VISp projections (**Fig. 4a**). Across the six major higher visual areas that Han et al. mapped (AM [VISam in our data], PM [VISpm], AL [VISal], LI [VISli], RL [VISrl], and LM [Visl]), we identified 7 enriched combinations and 6 depleted combinations. These combinations recapitulated all four under- or over-represented pairwise combinations reported by Han et al. In addition, we found three pairs (LM-AM, LI-AL, LI-AM) that were significantly enriched or depleted in our data; these were in the same direction as the trends reported by Han et al., which did not reach significance in that dataset. We then extended the pairwise combination analysis to all 13 cortical target areas, and sorted the areas based on their locations around the VISp (**Fig. 4b**). We found that the dominating collateralization patterns were seen across pairs of adjacent areas, which form two off-diagonal lines (**Fig. 4b**). Weaker enrichment extended to some non-adjacent pairs, but it is ambiguous whether these enrichments correspond to the classic dorsal and ventral streams or other organizations. These results therefore replicate previous findings on collateralization, but the underlying organization cannot be resolved by inspection of the enrichment alone.

**Figure 4.**
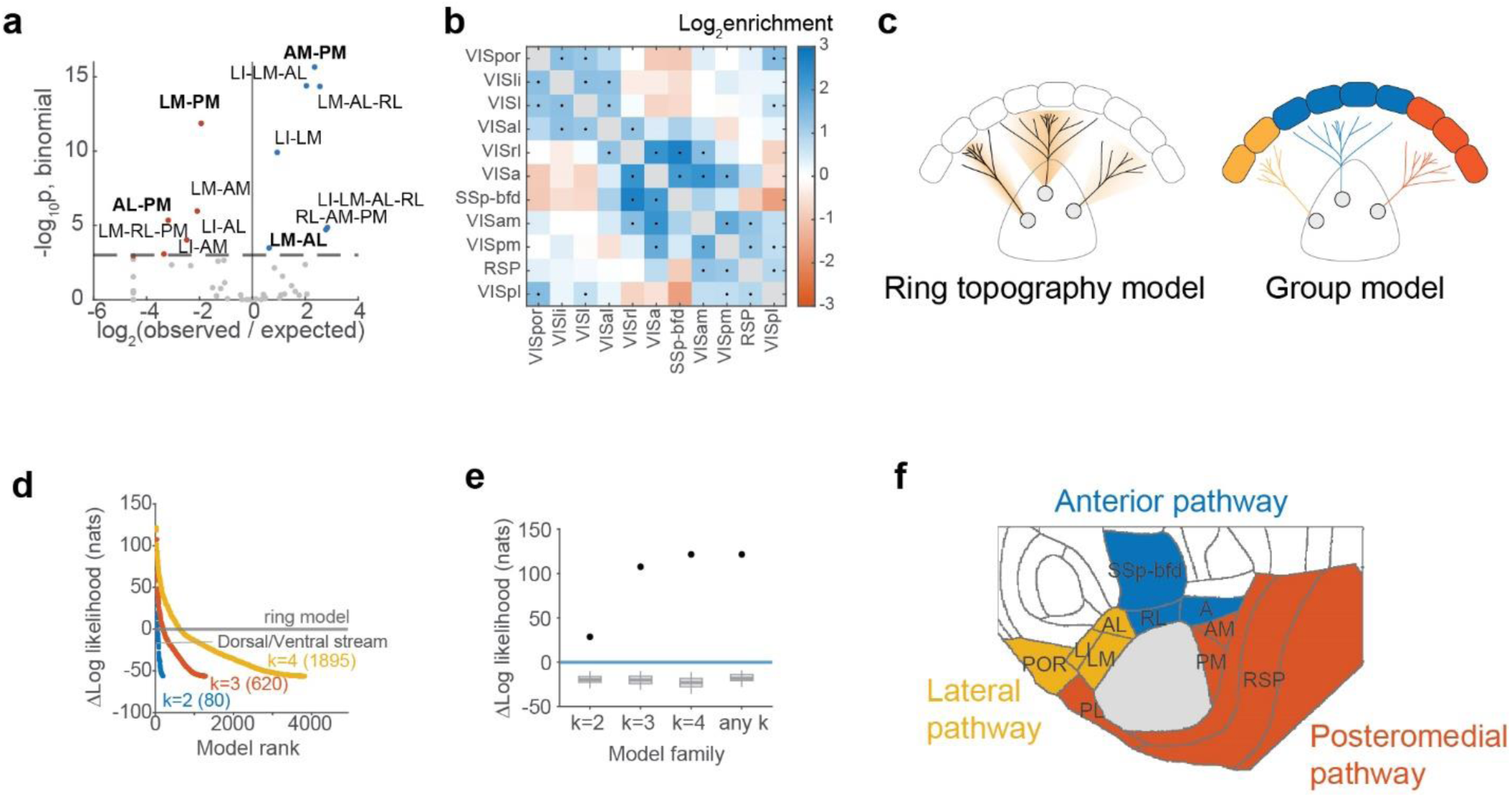
VISp neurons collateralize along three pathways. (**a**) Volcano plot showing the log odds ratio for collateral projection patterns, given individual projection probability. Horizontal dashed line indicates significance threshold (p < 0.05 Holm-Bonferroni correction). Significant combinations are labeled. Enriched combinations are shown in blue dots, and depleted combinations are red. Labels in bold indicate those that recapitulated Han et al., 2018. (**b**) Log enrichment for pairwise collateralization patterns, relative to independent individual projections. Significance is indicated by a black dot (p < 0.05, Holm-Bonferroni correction). (**c**) In the ring-topography model, each neuron preferentially projects to a contiguous range of areas around VISp, with projection probability decreasing with angular distance from its preferred direction. In the group model, areas are divided into discrete groups, and neurons are more likely to project to multiple areas within the same group than to areas across different groups. The strengths of within-group and across-group co-projection are fitted across all neurons. (**d**)The Δlog likelihood (y axis) of each group model compared to the ring model. Colors indicate model family based on group numbers. Within each group, the models are sorted from best performing to worst. The dorsal/ventral stream 2-group model is indicated (**e**) The y-axis shows the improvement in log-likelihood of the best-fitting group model relative to the ring-topography model. The dots indicate the best-fitting grouping observed in the data. Grey box plots show the null distribution of the best loglikelihood in each model family, obtained from 200 datasets simulated under the fitted ring-topography model and analyzed with the same grouping scan. Boxes indicate the interquartile range, center lines the median, and whiskers the 5th–95th percentiles. For all comparisons p = 0.005 (at the floor with 200 bootstraps). (**f**) The best grouping under 3-group models.

The strong enrichment of collateralization across neighboring areas suggested three possible sources of this organization. First, the pattern could reflect retinotopy, because different higher visual areas could represent partially overlapping or adjacent portions of visual space. Second, collateralization could follow a continuous topographic organization around the ring of higher visual areas surrounding VISp, such that nearby areas are preferentially co-targeted. Third, the apparent ring-like pattern could instead reflect discrete groups of areas that are spatially contiguous along the ring and preferentially collateralized as distinct pathways. We first tested the retinotopic explanation by comparing collateralization enrichment with and without accounting for the dependence of projection probability on retinotopic position (**ED Fig. 4a, b**). Accounting for retinotopy only marginally reduced the enrichment of collateralization patterns, indicating that retinotopic target biases were insufficient to explain the observed collateralization structure. We therefore next asked whether the remaining structure was better explained by a continuous ring-like organization or by discrete groupings of cortical targets into two, three, or four groups (**Fig. 4c**). Briefly, for each grouping, we fitted the binarized projection patterns with a pairwise maximum-entropy model, in which the pairwise association is determined only by whether a pair of projection targets are within the same group or not. For the ring model, we fitted an analogous model in which pairwise association depended instead on the angular distance between the two areas around VISp. Importantly, because the groupings only contribute to group membership of pairwise areas, all grouping models have the same number of free parameters as the ring model and thus matched in model complexity. We then compared the log-likelihoods across all candidate models and calibrated the best-performing model for each number of groups using a parametric bootstrap under the fitted ring model. Consistent with the observation that the enrichment of collateralization patterns does not follow the classic dorsal and ventral streams, the ring model outperformed a two-group model that separated the dorsal and ventral stream areas (**Fig. 4d**). However, the best performing model in each grouping number outperformed the ring model (**Fig. 4d, e**; p = 0.005 for all best-in-family models compared to the ring model, p value limited by permutation floor). The performance of the best models plateaus at three groups: VISpor, VISli, VISlm, and VISal as the lateral group, VISpm, VISam, RSP areas, and VISpl as the posteromedial group, and SSp-bfd, VISa, and VISrl as the anterior group (**Fig. 4f**). Relative to the retinotopy model, the ring model gained 259 nats and the optimal three-group model gained 367 nats, placing the three-group improvement well beyond what ring-like organization alone accounts for. The grouping was also largely stable: the three best models differed only in the assignment of VISal and VISa, which lie near group boundaries, and the next two models performed only modestly worse than the optimum (Δlog-likelihood relative to the ring model 92.9 and 85.6 nats, compared with 107.5 nats for the best model; **ED Fig. 4c**). Thus, rather than recovering the canonical dorsal-ventral division, our data resolved three preferential collateralization pathways – lateral, posteromedial, and anterior. Notably, the posteromedial and anterior pathways separate targets that have often been grouped together within the dorsal visual stream, consistent with previous anatomical and functional evidence that the mouse dorsal stream contains distinct substreams (D’Souza et al., 2022; Han et al., 2022; S. Yao et al., 2023).

### Axonal BARseq2 resolves transcriptomic identities of neurons in the visual cortex

Our analyses so far showed that VISp projection patterns are structured by both retinotopic position and specific collateralization patterns. We next asked whether and how transcriptomic identity provides an additional dimension of organization for these projections. Because axonal BARseq2 measures endogenous gene expression and projections in the same neurons, it allowed us to directly relate transcriptomic identity with projection patterns at single-cell resolution. We first used hierarchical clustering to define transcriptomically defined cell types (**Fig. 5a, b**) across pooled barcoded and non-barcoded neurons after QC (see Methods), resulting in two levels of clusters that largely correspond to subclass level and supertype/cluster level in the Allen Brain Cell (ABC) atlas (Z. Yao et al., 2023). In the cortex, clustering resulted in subclasses that corresponded to all major cortical excitatory neuron types, including four subclasses of IT neurons, L5 ET, L6 CT, L6b, NP, and Car3. These subclasses showed the expected laminar organization in VISp (**Fig. 5c**). Additional area-specific excitatory subclasses were seen in the retrosplenial cortex and the entorhinal cortex (**Fig. 5a, b**). Because these cell types were not found in large numbers in the visual cortex, we excluded them from subsequent analyses. Neurons from the two brains were intermingled in transcriptomic UMAP space (**ED Fig. 5a**), and each major subclass and transcriptomic cluster contained a substantial number of cells from both brains, indicating that the identified cell types were reproducible across animals (**ED Fig. 5b, c**).

**Figure 5.**
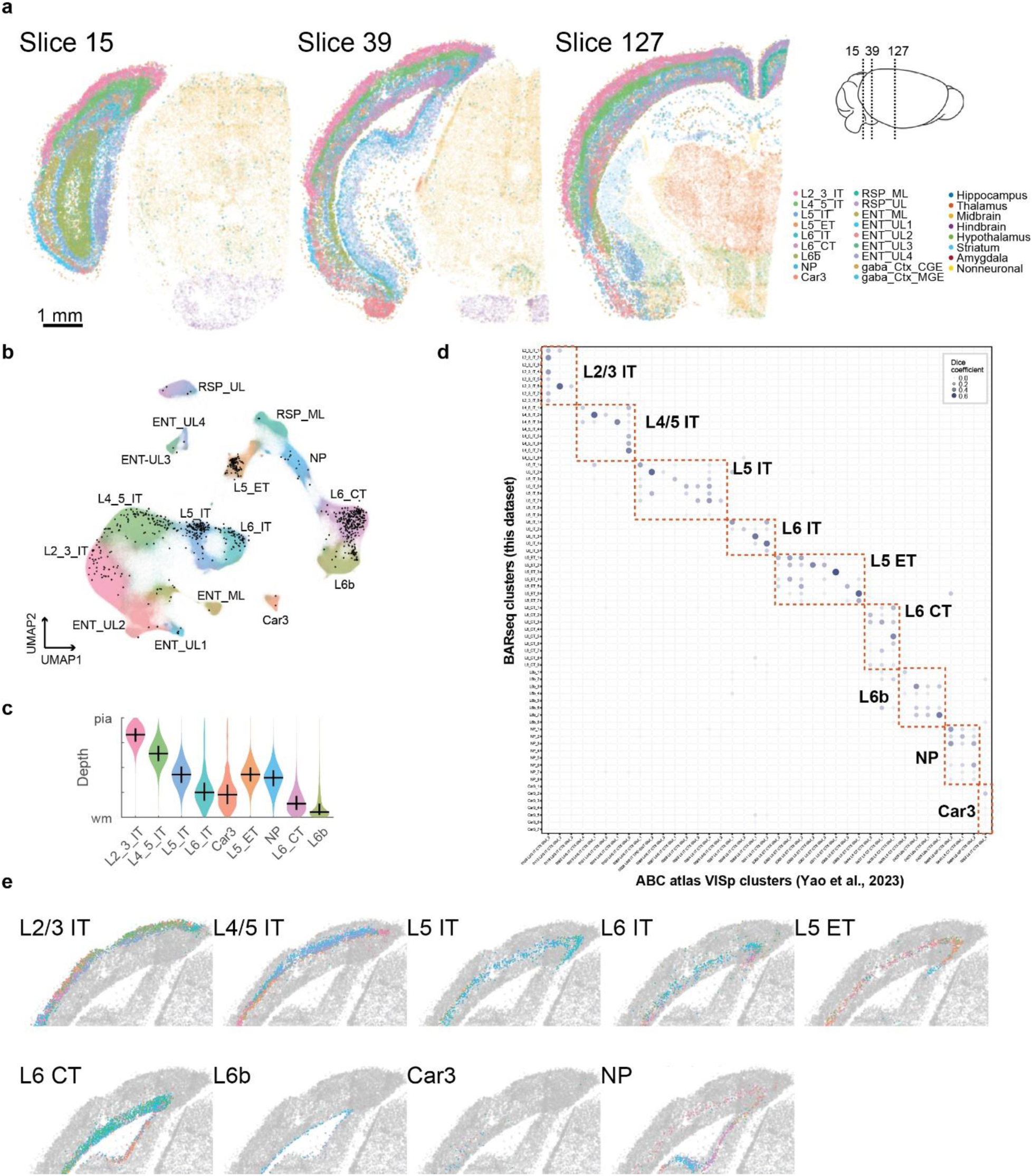
Axonal BARseq2 reveals gene expression of projection neurons. (**a**) Images from three representative slices, showing cells colored by subclasses. The locations of the three sections are shown on the model brain on the right. (**b**) UMAP plot of cortical excitatory neurons, colored by subclasses. (**c**) Laminar distribution of the nine cortical excitatory subclasses that are in the VISp. (**d**) Overlap (dice coefficient) between BARseq cell types (this dataset) and ABC atlas (Z. Yao et al., 2023) clusters in VISp. Dashed boxes indicate subclasses. (**e**) Spatial distribution of BARseq cell types within each subclass. Cell types from the same subclass are plotted together and shown in different colors.

To determine whether our data recapitulated previously defined transcriptomic cell types, we mapped individual neurons to reference single-nucleus RNA-seq data from VISp in the ABC Atlas (Z. Yao et al., 2023) using a k-nearest neighbor based approach (Chen et al., 2025), and assessed overlap between our cluster labeling and the reference taxonomy (**Fig. 5d**; see **ED Fig. 5d** for mapping of all isocortical cell types). Each BARseq cluster mapped onto a small number of reference clusters, suggesting that our dataset recapitulated transcriptomic diversity across reference transcriptomic datasets (**Fig. 5d**). Consistent with previous studies(Chen et al., 2025; Z. Yao et al., 2023; Zhang et al., 2023), fine-grained cell types within each subclass were differentially enriched across cortical areas and/or sublaminae (**Fig. 5e**). These results indicate that our data recapitulates both the molecular and spatial organization of previously defined cell types.

### Fine-grained cell types distinguish corticocortical projections to the three pathways

Having established that our data recapitulate known transcriptomic cell identities, we next examined how their transcriptomic identities relate to their projection patterns. At the subclass level, neurons projected to targets that were consistent with their transcriptomic identities (**Fig. 6a**). The four IT subclasses largely projected to the cortex, L5 ET neurons projected to the superior colliculus, and L6 CT neurons projected to the thalamus.

**Figure 6.**
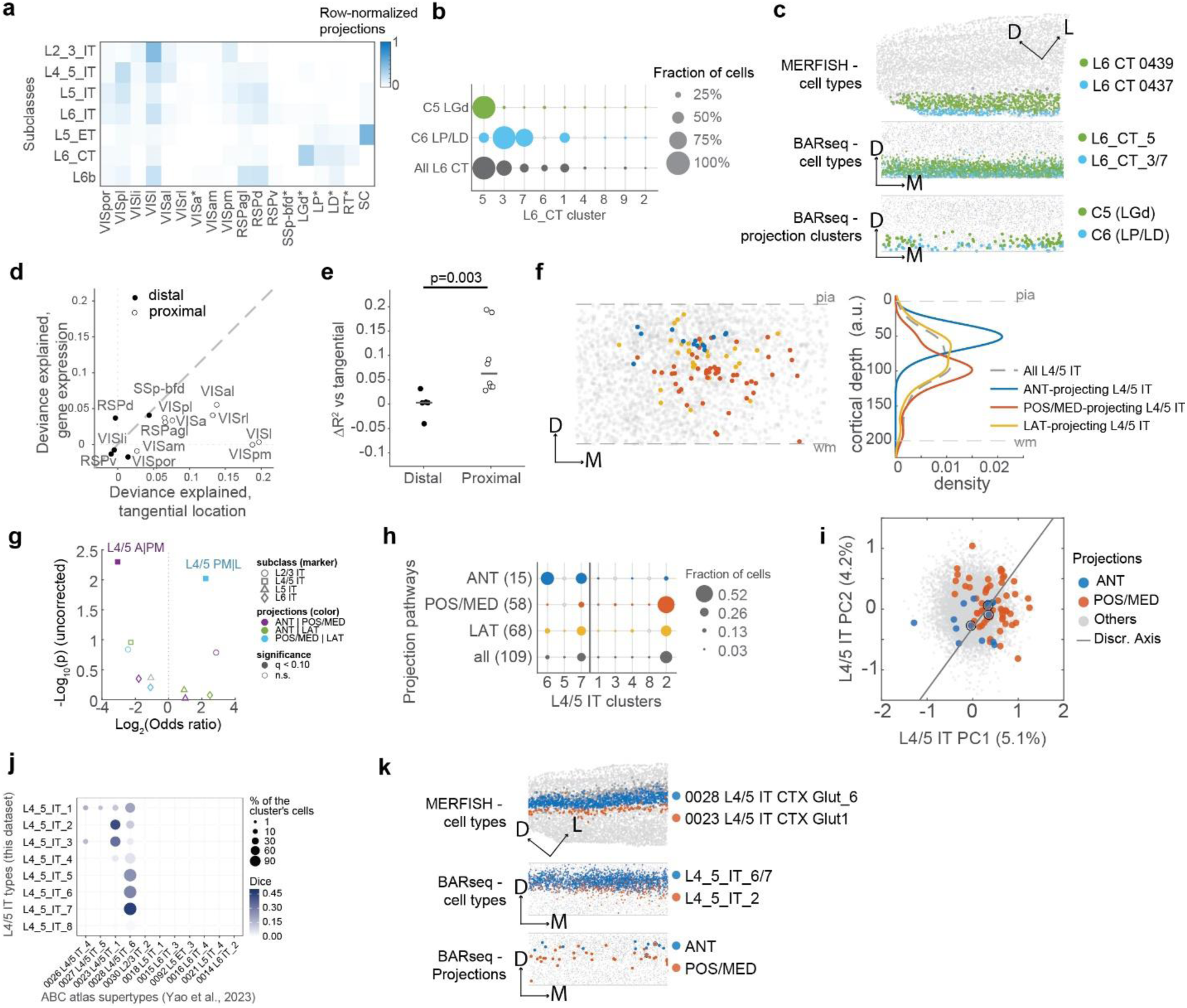
Transcriptomically defined L4/5 IT types segregate VISp output to visual streams. (**a**) Pseudobulk projection patterns for neurons in each subclass. Only subclasses with at least 10 barcoded neurons are shown. (**b**) The fraction of neurons from each projection cluster that belongs to each L6 CT transcriptomic type. The transcriptomic types are sorted by abundance across barcoded neurons. (**c**) A “coronal” view of neurons of the indicated categories, including ABC atlas clusters (*top*), BARseq transcriptomic types (*middle*), and BARseq projection clusters (*bottom*). The ABC atlas MERFISH data is shown in a tilted coronal view of a single section, whereas the BARseq data are stacked sections shown in flatmap coordinates, where the x axes indicate the medial-lateral axis, and the y axes indicate depth. (**d**) Cross-validated deviance in projections to each area, explained by tangential locations (x axis) and by gene expression (y axis). Distal areas are shown by solid dots, and proximal areas are shown by circles. (**e**) ΔDeviance explained between gene expression compared to tangential locations for distal areas and proximal areas. Horizontal bars indicate mean. p = 0.003 using rank-sum test. (**f**) L4/5 IT neurons from both brains that project to each of the three pathways, plotted on a flatmap coronal view (x axis showing medial-lateral axis, and y axis showing depth). A smoothed histogram on the right shows the laminar distribution profile for the three projection-defined neuronal populations, and all L4/5 IT neurons. (**g**) Transcriptomic segregation of L4/5 IT projection groups. Each point compares two projection groups across pseudotime-ordered transcriptomic clusters. The x-axis shows the log odds ratio at the partition that best separates the groups; the y-axis shows significance from a position-matched permutation test that also accounts for partition selection. Colors indicate projection-group pairs. Filled symbols indicate FDR < 0.1. (**h**) The fraction of neurons that project to each pathway and belong to each L4/5 IT transcriptomic types (row normalized to 1). The transcriptomic types are sorted by their mean values on a pseudotime axis defined across all IT neurons. The vertical line indicates the transcriptomic partition that best separated projection-group preferences, as used in (g). (**i**) Scatter plot showing L4/5 IT neurons on the first two principal components derived from all L4/5 IT neurons in VISp. Dot colors indicate projection pathways, and the solid line indicates the discriminant axis between the two projection populations. (**j**) Overlap (dice coefficient) between BARseq transcriptomic L4/5 IT types and ABC atlas L4/5 IT supertypes. Colors indicate dice coefficient and circle size indicates fraction of cells in the BARseq transcriptomic type. (**k**) A “coronal” view of neurons of the indicated categories, including ABC atlas clusters (*top*), BARseq transcriptomic types (*middle*), and BARseq projection clusters (*bottom*). The ABC atlas MERFISH data is shown in a tilted coronal view of a single section, whereas the BARseq data are stacked sections shown in flatmap coordinates, where the x axes indicate the medial-lateral axis, and the y axes indicate depth.

We next asked whether this correspondence extended beyond broad transcriptomic subclasses to finer-grained cell types. We focused on the two projection-defined L6 CT populations, C5 and C6, which projected predominantly to LGd and LP/LD, respectively. These populations were associated with distinct transcriptomic types: C5 LGd-projecting neurons were mostly L6_CT_5, whereas C6 LP/LD-projecting neurons were enriched for L6_CT_3 and L6_CT_7 (**Fig. 6b**). We then asked whether these transcriptomic assignments were also consistent with the laminar positions of the two projection-defined populations. All four L6 CT types mapped to the ABC Atlas supertype “0114 L6 CT CTX Glut_1”; at the finest cluster level, L6_CT_3/7 mapped predominantly to cluster 0437, whereas L6_CT_5 mapped predominantly to cluster 0439 and only weakly to 0437 (**Fig. 5d**). In the ABC Atlas MERFISH data, cluster 0437 was concentrated in deeper L6, whereas cluster 0439 occupied a more superficial position (**Fig. 6c**). This laminar organization mirrored that of the projection-defined populations, with C5 LGd-projecting neurons located more superficially than C6 LP/LD-projecting neurons. Thus, the relationship between transcriptomic identity and projection pattern extends from broad IT, ET, and CT subclasses to fine-grained L6 CT types, demonstrating that our dataset can distinguish transcriptomic differences among projection-defined populations at single-cell resolution.

For IT neurons, projections to areas across the three pathways appeared to be differentially distributed across layers (**ED Fig. 6**), but projections to many areas were shared across the subclasses (**Fig. 6a**). This relatively broad overlap contrasted with the strong dependence of projection probability on the tangential position of neurons within VISp (**Fig. 3h**). To compare the contributions of spatial position and gene expression to target selection, we fitted ridge-regularized logistic regression models to predict whether individual neurons projected to each target area using either their tangential soma coordinates or the top 15 principal components of gene expression (**Fig. 6d**). Tangential position explained up to ∼20% of the variance in projection probability, whereas the top 15 gene-expression PCs explained at most ∼5%. Consistent with the stronger spatial biases observed for areas proximal to VISp, the advantage of tangential position over gene expression was substantially greater for proximal than for distal targets (**Fig. 6e**; additional variance explained by tangential location is 0.09 ± 0.07 for proximal areas and 0.00 ± 0.03 for distal areas, mean ± s.d., p = 0.003 rank-sum test). Moreover, for many proximal targets, these spatial biases were preserved across transcriptomic subclasses. For example, VISpm-projecting neurons were consistently enriched medially within VISp, whereas VISl-projecting neurons were enriched laterally, across multiple subclasses (**ED Fig. 5e**). Together, these results indicate that projection variability is more strongly associated with the retinotopic position of the soma than with broad transcriptomic variation, particularly for proximal targets.

The relatively weak relationship between broad gene-expression variation and projection specificity raised the possibility that projection biases emerge at finer transcriptomic resolution (Kim et al., 2020). Consistent with finer-scale projection organization, neurons projecting to the three pathways were differentially distributed across cortical depth even within the narrow laminar band occupied by L4/5 IT neurons (**Fig. 6f**). Neurons projecting to the anterior, lateral, and posteromedial pathways were preferentially located toward the upper, middle, and lower portions of this band, respectively. Because transcriptomically defined cell types often occupy distinct sublaminar positions, these laminar differences suggested that fine-grained transcriptomic types within a subclass might also differ in their pathway preferences.

To test this possibility, we partitioned the fine-grained clusters within each subclass into two transcriptomically contiguous groups and identified the partition that best separated each pair of pathways (see **Methods**). In L4/5 IT neurons, the three pathways showed significant segregation across these transcriptomic partitions (**Fig. 6g**; FDR < 0.1 between the anterior pathway and the posteromedial pathway, and between the lateral pathway and the posteromedial pathway, based on a permutation test that accounted for both optimization over transcriptomic cut points and tangential spatial biases in projection probability): The posteromedial projections were enriched in L4/5_IT_2, the anterior projections were highly enriched in clusters L4/5_IT_6 and 7, and the lateral projections were distributed across the two groups (**Fig. 6h**). This projection pathway preference was also captured by low-dimensional transcriptomic variation within L4/5 IT subclass: The first two principal components of L4/5 IT subclass could both distinguish the anterior and posteromedial projection populations (PC1 AUROC 0.72, p =0.005; PC2 AUROC 0.70, p = 0.01, using permutation tests), and a classifier based on the two PCs achieved an AUROC of 0.78 (p = 0.002 using a permutation test; p = 0.006 using a permutation null that was additionally matched on tangential soma positions; **Fig. 6i**). In contrast, comparable molecular partitions did not distinguish the three pathways within other subclasses (**Fig. 6g**), although our sampling was sparse in L2/3 IT (n = 43 neurons). Thus, the pathway-associated differences were evident both across fine-grained transcriptomic clusters and along major axes of gene-expression variation.

We next asked whether the transcriptomic differences associated with these pathways were consistent with their laminar organization in an independent reference dataset. In the ABC Atlas, L4/5_IT_6 and L4/5_IT_7, which were enriched among anterior-projecting neurons, mapped predominantly to supertype 0028, whereas L4/5_IT_2, which was enriched among posteromedial-projecting neurons, mapped predominantly to supertype 0023 (**Fig. 6j**). MERFISH data from the ABC Atlas showed that these reference supertypes occupied distinct laminar positions that matched the relative cortical depths of the corresponding projection-defined populations in our dataset (**Fig. 6k**). In particular, the supertype associated with posteromedial-projecting neurons occupied a deeper position than the supertype associated with anterior-projecting neurons, consistent with the depth-dependent pathway bias observed in our data (**Fig. 6f**). Together, the cluster-level associations, their expression along low-dimensional transcriptomic axes, and their matching laminar organization in the ABC Atlas provide convergent evidence that collateralization pathway preference in L4/5 IT neurons is associated with fine-scale transcriptomic organization.

## Discussion

We used axonal BARseq2 to simultaneously measure gene expression and axonal projections from individual VISp neurons to assess the contribution of their topographical locations and transcriptomic identities on projection patterns. We found that retinotopy predominantly determines projection probability to individual cortical targets, particularly for areas proximal to VISp, but cannot explain which targets are co-innervated. Instead, collateralization patterns can be divided into three cortical pathways that broadly correspond to the ventral stream and two subdivisions of the dorsal stream. These pathways are further associated with fine-grained transcriptional identities within L4/5 IT neurons, providing anatomical evidence that visual-stream segregation is anatomically instantiated by distinct transcriptomically defined neuronal populations. Together, our results suggest that VISp output is organized along separable axes: retinotopic position constrains where individual projections are sent, whereas cell-type-associated collateralization determines how targets are coupled into visual processing pathways.

### Axonal BARseq2 reveals distinct wiring rules for proximal and distal VISp projections

Topography and cell identity are major determinants of long-range projection patterns, yet few approaches can resolve both in the same neurons at sufficient scale to determine how they differentially contribute. By combining axonal BARseq with BARseq2, axonal BARseq2 jointly measures soma position, transcriptomic identity and projections. We extensively validated both the transcriptomic and anatomical resolution of the resulting measurements against independent single-cell transcriptomic, single-neuron tracing and known retinotopic datasets. Axonal BARseq2 therefore complements single-neuron reconstruction approaches: it sacrifices some spatial resolution in exchange for substantially denser sampling and the ability to link projection patterns directly to gene expression in situ. This combination makes it particularly well suited for resolving the joint spatial and molecular organization of long-range projections, including in systems where registration across individuals is challenging.

Resolving these two sources of organization is particularly important in VISp, where both tangential position and cell type identity have been associated with projection target choice (Sorensen et al., 2026). Across nearly the full extent of VISp, we found that tangential position was substantially more predictive of corticocortical target choice than broad transcriptomic variation, particularly for targets proximal to VISp. Moreover, these spatial biases closely followed the retinotopic organization of VISp and its higher visual area targets, suggesting that much of the previously described dependence of projection probability on cortical position reflects preservation of visual-field organization rather than a generic effect of soma location. Strikingly, this dependence was concentrated among proximal targets and was much weaker for distal cortical areas. These results suggest that VISp output may be governed by distinct spatial constraints at different projection target distances: retinotopy strongly constrains routing to nearby cortical targets, whereas projections to more distant networks are less tightly coupled to the neuron’s position in the visual map. This proximal-distal distinction refines the emerging view that spatial position and molecular identity jointly constrain cortical connectivity and raises the possibility that differential topographic constraints on local and long-range projections represent a more general wiring principle across cortex.

### Selective collateralization segregates VISp output into visual processing streams

Single-neuron tracing has challenged a simple labelled-line model of corticocortical communication. Han et al. showed that most VISp neurons project to multiple cortical targets, often in non-random combinations, suggesting that individual neurons broadcast signals across select subsets of areas (Han et al., 2018). However, these collateralization patterns did not align clearly with the canonical division of higher visual areas into dorsal and ventral streams, leaving it unclear whether collateralization contributes to stream segregation or instead reflects other features of cortical organization. We recapitulated the projection combinations identified by Han et al. and, by extending this analysis across a broader set of cortical targets and explicitly accounting for retinotopic biases, found that collateralization could not be explained by either retinotopy or the continuous topographic arrangement of areas around VISp. Instead, co-projection patterns were better described by three discrete pathways. Thus, the broad divergence of VISp projections need not imply indiscriminate mixing of information across cortical networks. Rather, visual streams may be instantiated through *selective broadcasting*: individual neurons distribute signals to multiple cortical areas, while structured collateralization constrains which areas receive those signals together.

This organization also supports the view that the dorsal stream is better described by multiple parallel pathways than by a single stream. The classical framework divided visual processing into ventral and dorsal pathways associated broadly with object identification and spatial processing (Mishkin et al., 1983; Mishkin and Ungerleider, 1982), with the dorsal stream subsequently reframed around visually guided action (Goodale and Milner, 1992). Kravitz et al. (Kravitz et al., 2011)further argued that the dorsal stream is not a single pathway, but branches into distinct circuits supporting spatial working memory, visually guided action and navigation, and more recent anatomical and functional studies in mice have similarly found substantial specialization within areas conventionally assigned to the dorsal stream (D’Souza et al., 2022; Han et al., 2022). Our collateralization-defined pathways correspond to this finer functional organization. The anterior pathway preferentially groups visual areas associated with sensorimotor processing, including VISrl and VISa, together with somatosensory cortex, whereas the posteromedial pathway groups VISpm and VISam with retrosplenial cortex, a network strongly implicated in spatial orientation and navigation. Thus, although the cortical areas and circuit architecture differ between rodents and primates, the separation we observe broadly parallels the distinction between action-related and posteromedial navigation-related branches proposed for the primate dorsal stream. Importantly, whereas these pathways have generally been defined by specialization and connectivity downstream of early visual cortex, our results show that their segregation is already apparent in the collateralization patterns of individual VISp neurons. This suggests that diversification of the dorsal stream may begin at the single-neuron output level of the primary visual cortex rather than downstream.

### Stream segregation by transcriptomic identity

The selective association of the visual pathways with fine-grained cell types in L4/5 IT suggests that this population may play a specialized role in separating visual cortical output into distinct processing streams. Within the relatively narrow L4/5 IT band, anterior- and posteromedial-projecting neurons were distinguished by both transcriptomic identity and laminar position, whereas comparable molecular divisions were not apparent in the other IT subclasses. The lack of effect in L2/3 IT neurons does not conflict with prior evidence for projection-specific gene expression in this population (Kim et al., 2020), because our sampling of L2/3 IT subclass was sparse and underpowered. Furthermore, lateral- and anterior-projecting L2/3 neurons in mouse VISp are separated along a continuous transcriptomic axis that may not be captured by transcriptomically defined types (Kim et al., 2020). Whether stream segregation is specific to fine-grained types of L4/5 IT neurons or extends more broadly across superficial IT populations will require future study with denser sampling and measurement of axonal collateralization.

This organization is particularly intriguing in light of primate V1: The mouse L4/5 IT neuron is transcriptomically homologous to the primate L4 IT neurons(Bakken et al., 2021; BRAIN Initiative Cell Census Network (BICCN) et al., 2021; Hodge et al., 2019; Z. Yao et al., 2023), which are the predominant population in the primate L4A/B/C, and layer 4B forms a major source of corticocortical output and contains specialized neuronal populations that differentially innervate extrastriate pathways, including V2, MT, and V6 (Galletti et al., 2001; Nassi and Callaway, 2007; Sincich and Horton, 2003; Yarch et al., 2019). Thus, although the precise cell types and laminar architecture differ substantially between rodents and primates, our results raise the possibility that assigning distinct visual output pathways to fine-grained neuronal identities within the L4/5 IT subclass represents a conserved organizational principle. We propose that this may be a conserved property of the L4/5 IT population, despite extensive primate-specific diversification and specialization of neurons in L3-L4, including primate-specific transcriptomic populations. Such a conserved wiring rule could be retained even as primate visual streams become more elaborate through the addition of intermediate cortical areas and expansion and specialization of cortical cell types.

Conserved stream segregation in the L4/5 IT population could nonetheless underlie substantially elaborated downstream pathways across species. The mouse anterior pathway provides one example. VISrl groups with somatosensory cortex in our collateralization analysis, consistent with evidence that VISrl integrates topographically corresponding visual and somatosensory signals and contributes to a multimodal representation of near-body space (Zhang et al., 2026). These properties resemble those of the primate ventral intraparietal area (VIP), whose neurons integrate visual and somatosensory information in representations of near space (Avillac et al., 2007, 2005; Duhamel et al., 1998). Yet, unlike mouse VISrl, primate VIP is not known to receive substantial direct input from V1; visual signals instead reach VIP through intervening dorsal-stream areas, including MT and V6 (Galletti et al., 2001; Maunsell and van Essen, 1983). The mouse anterior pathway may therefore represent a more compact version of a circuit that was expanded in primates by inserting additional target regions between the primary visual cortex and a functionally related parietal network. The posteromedial pathway suggests a complementary form of elaboration. In mice, VISp neurons in this pathway directly collateralize to retrosplenial cortex together with posteromedial visual areas. In contrast, anatomical tracing in macaques has found little or no direct V1 input to retrosplenial cortex, which instead receives visual cortical input through extrastriate areas including V2 and area 19 (Kobayashi and Amaral, 2003; Morris et al., 1999). Thus, the anterior and posteromedial pathways illustrate two ways in which a common functional architecture could be modified across evolution: a processing stream can be extended through the addition of intermediate cortical stages, or a direct connection can be replaced by routing through intervening visual areas.

## Supplementary Notes

### Supplementary Note 1: Barcode matching and quality control

To assess whether barcodes in axons and somata could be matched unambiguously, we calculated the minimum Hamming distance from each axonal barcode to the full set of soma barcodes and, as a control, to a shuffled set of soma barcodes. 1,438,249 barcode rolonies fell within one mismatch of the 10,616 soma barcodes, compared with only 26,830 within one mismatch of shuffled soma barcodes (**ED Fig. 2a**), indicating that 1 – 26,830/1,438,249 = 98.1% of barcode rolonies were matched to the correct soma. To further reduce the impact of chance matches, we leveraged the spatial continuity of real anatomical structures, such as axons, to distinguish them from spurious ones. We reasoned that a barcode rolony located near other rolonies matched to the same neuron is more likely to represent a real match than an isolated rolony. We therefore used DBSCAN(Ester et al., 1996) to spatially cluster the rolonies matched to each neuron and removed any rolonies that did not cluster with others.

The impact of chance matches on individual neurons, however, depends on how many rolonies each neuron matches: two spurious rolonies negligibly affect the mapping accuracy of a neuron with 100 matching rolonies, but would render a neuron with only five matching rolonies effectively uninterpretable. We reasoned that chance matches should be random and thus spread across many distinct barcodes, whereas true matches should converge on a single barcode. Indeed, most neurons matched relatively few unique barcodes given their total number of matched rolonies, whereas a small subset matched many distinct barcodes at one mismatch (**ED Fig. 2b**) and had correspondingly few rolonies for the primary barcode (the matched barcode with the most rolonies) (**ED Fig. 2c**). We therefore excluded all neurons for which the ratio of unique matched barcodes to matched barcode rolonies exceeded 0.4. Combined with additional quality-control filtering, these criteria resulted in 1,448 high-quality neurons for downstream analysis, each with a soma in VISp and at least five barcode rolonies outside VISp.

## ED Figure legends

**Extended Data Figure 1.**
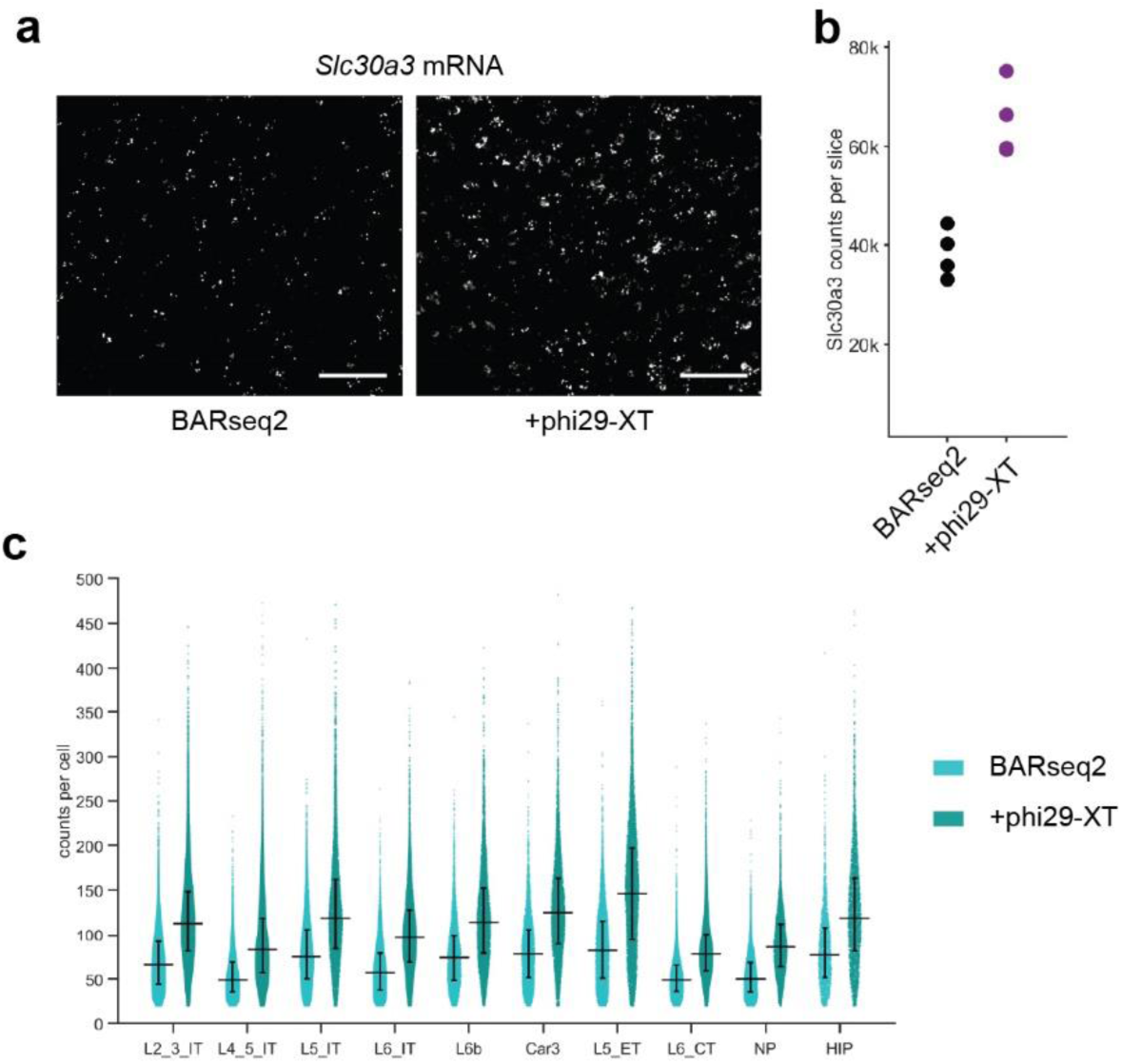
Optimization of axonal BARseq2. (**a**) Example images showing amplification and detection of *Slc30a3* using the original BARseq2 protocol (*left*) and with additional phi29-XT amplification (*right*). Scale bar = 50 µm. (**b**) *Slc30a3* counts across four sections using BARseq2 and with additional phi29-XT amplification. Each dot shows the total counts from one coronal section. (**c**)Violin plots showing the distribution of reads per cell in major isocortical excitatory neuron subclasses, sequenced using the 104-gene panel. Black bars indicate mean and standard deviations.

**Extended Data Figure 2.**
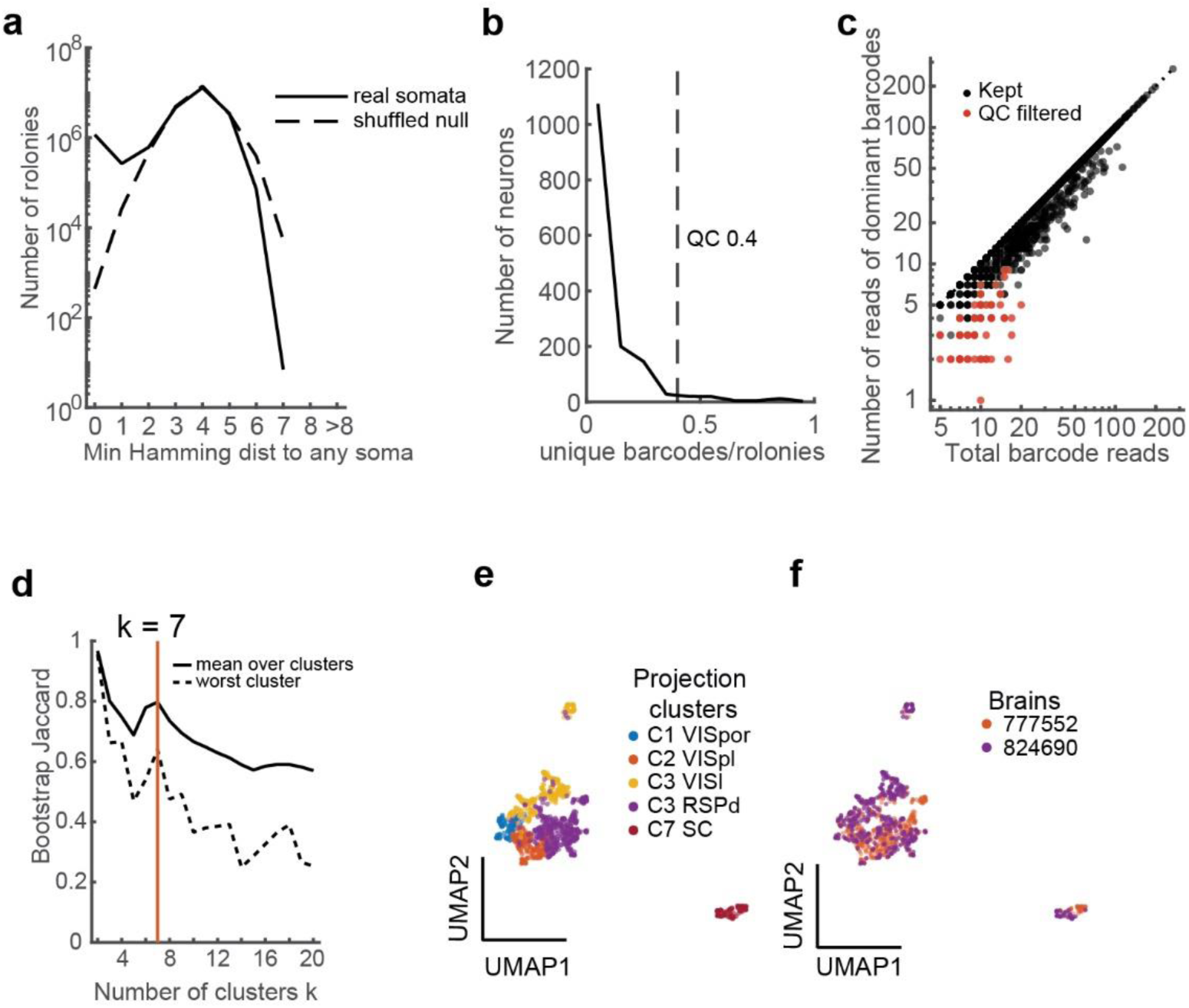
Barcode matching and quality control. (**a**) Histogram of minimal hamming distance from each barcode rolony to real barcodes in somas and to a shuffled control. (**b**) Distribution of neurons with the indicated ratio between unique barcodes recovered from matching axonal rolonies and the total number of rolonies. Dashed vertical line indicates quality control threshold. (**c**) Scatter plot showing total barcode reads (x axis) and the number of reads from the dominant barcode (y-axis). Red dots indicate neurons that are filtered out by the QC threshold in (b). (**d**) Clustering stability (Jaccard index across bootstraps) for the indicated number of projection clusters. Solid line shows the mean across clusters and dashed line indicates the least stable cluster. (**e**)(**f**) UMAP plots of projection neurons, colored by cluster (e) or by brain (f). Only IT and L5 ET neurons are shown, because the thalamus is largely not sampled in 824690 and L6 CT neurons are dominated by brain 777552.

**Extended Data Figure 3.**
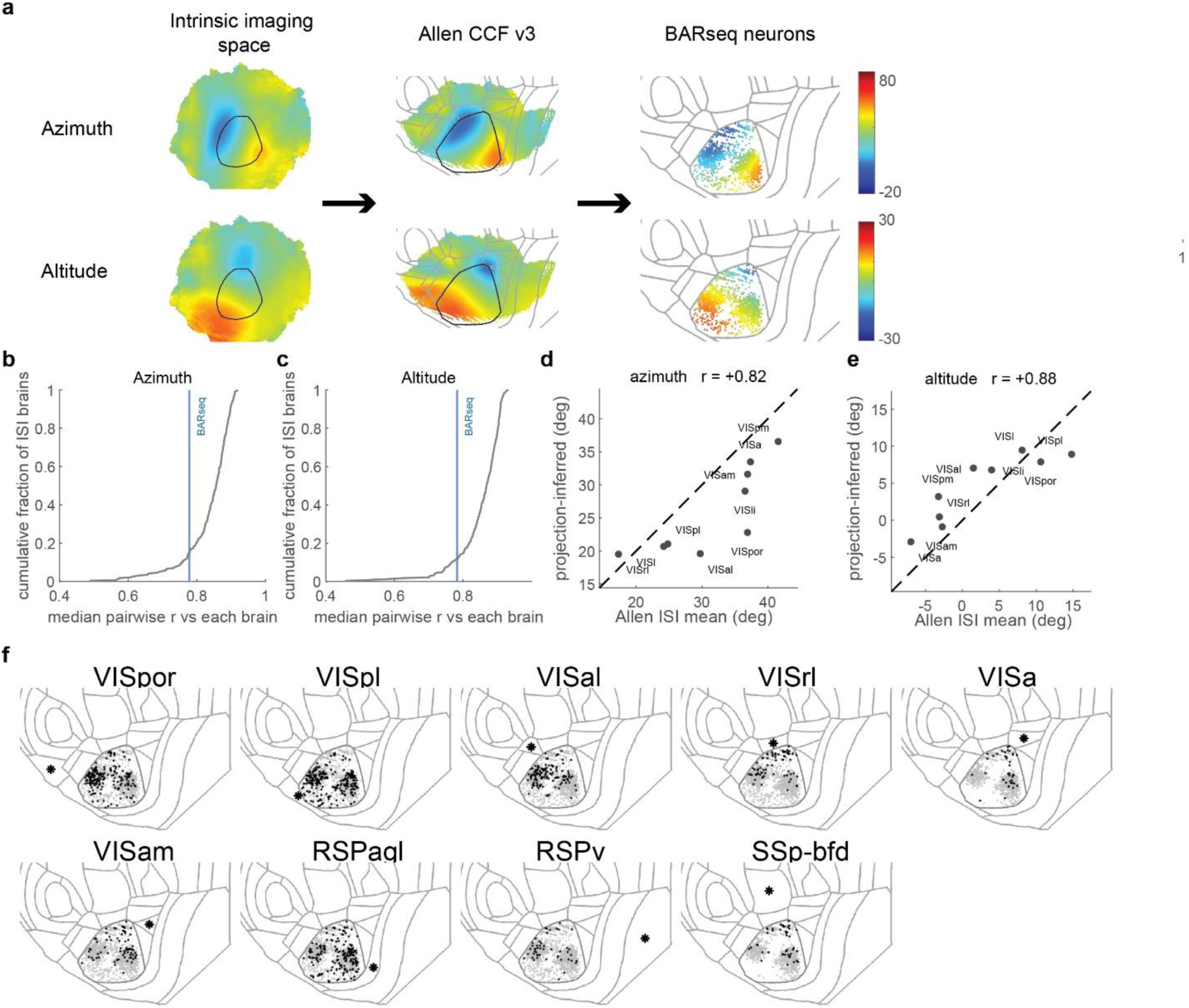
Axonal BARseq2 recapitulates retinotopy. (**a**) Intrinsic imaging-based retinotopic maps in the Allen mouse brain connectivity atlas (*left*) are first transformed to CCF space (*middle*), then the azimuth and altitude values are assigned to somas of neurons in this dataset (*right*). (**b**)(**c**) Cumulative distribution function of the pixel-wise correlation between the mean azimuth and altitude maps of each brain in the Allen mouse brain connectivity atlas to all other brains in the dataset. Blue vertical line indicates correlation between the mean azimuth and altitude maps estimated from this dataset to all brains in the Allen mouse brain connectivity atlas. (**d**)(**e**) The mean azimuth (d) and altitude (e) values for each cortical area inferred from the intrinsic imaging data (x axis) and our data (y axis). Pearson correlations are indicated on top. (**f**) The locations of neuronal somata (black dots) that project to the indicated target cortical areas. Neurons that did not project to the target areas are shown in gray, and the target areas are shown with an asterisk.

**Extended Data Figure 4.**
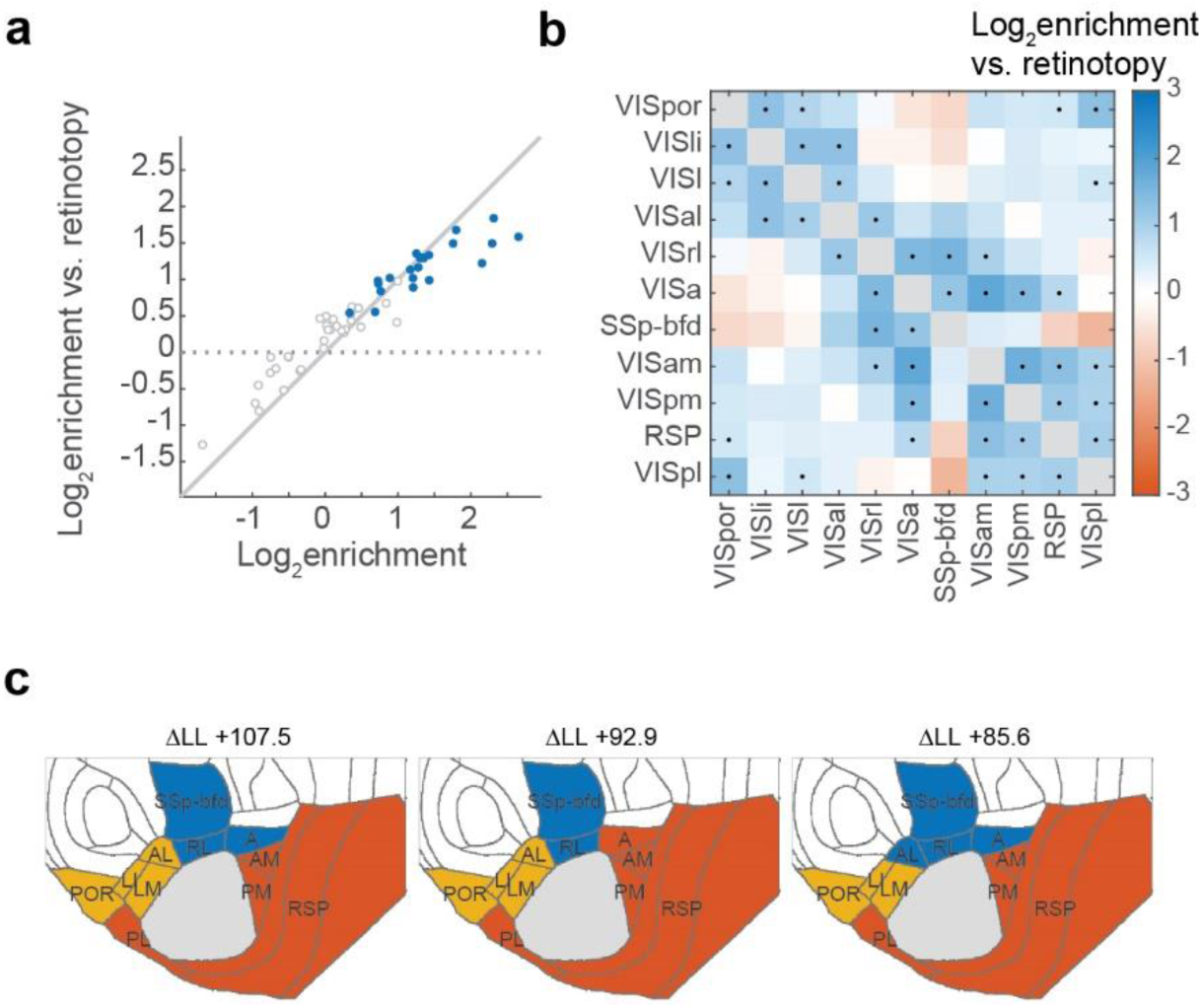
VISp neurons collateralize along three pathways. (**a**) Pairwise co-projection enrichment before and after accounting for retinotopy. The x-axis shows log enrichment relative to that expected from the marginal projection probabilities of the two targets, and the y-axis shows log enrichment relative to a retinotopy-conditional model, in which each neuron’s target-specific projection probabilities are predicted from its soma position and targets are otherwise independent. Blue points indicate target pairs that remain significantly enriched after accounting for retinotopy. The dotted horizontal line indicates enrichment fully explained by retinotopy, whereas the diagonal line indicates no effect of retinotopy. (**b**) Log odds ratio for pairwise collateralization patterns, after accounting for a retinotopy-conditional model. Significance is indicated by a black dot (p < 0.05, Holm-Bonferroni correction). (**c**) Top 3 groupings under the 3-group models. The improvements in log likelihood are indicated on top.

**Extended Data Figure 5.**
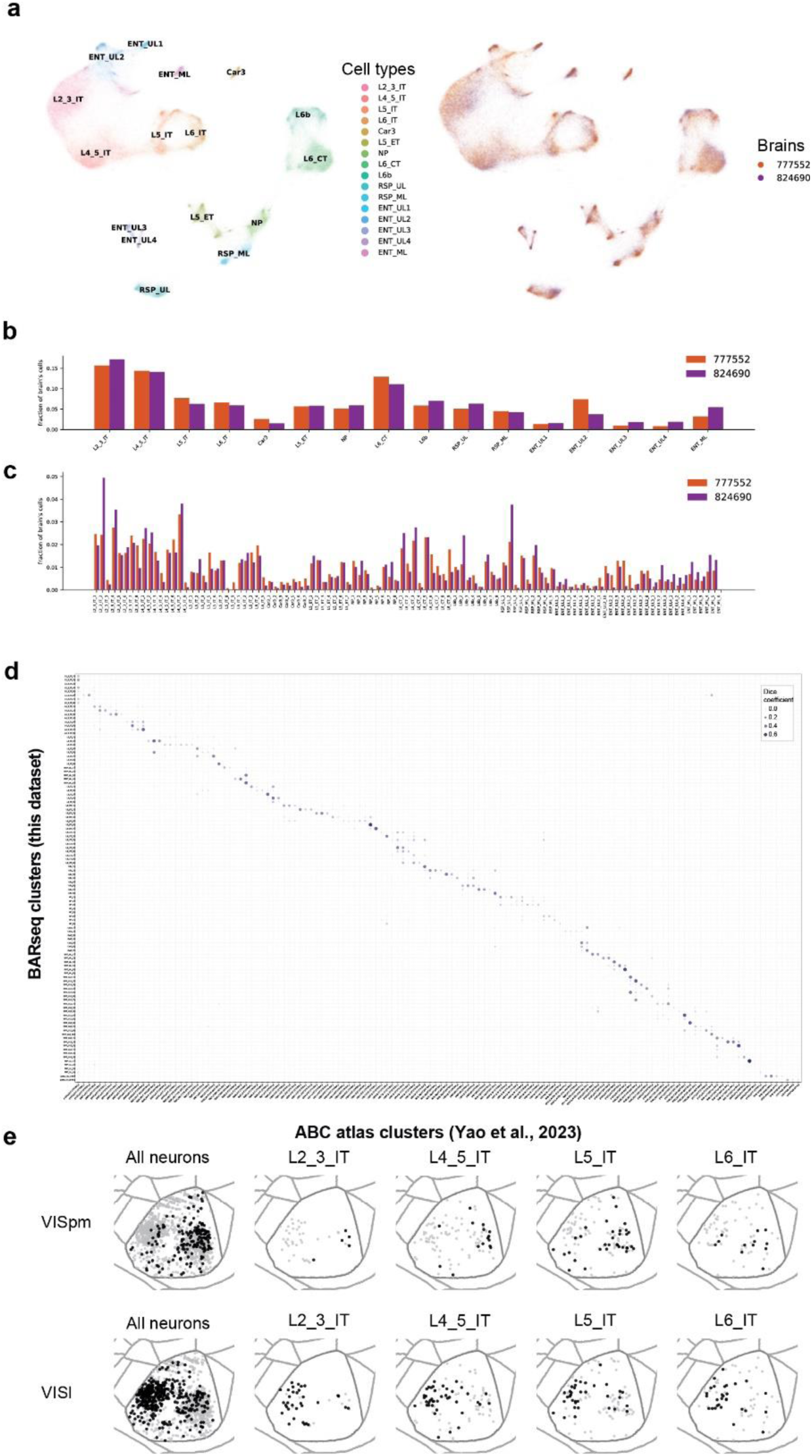
Axonal BARseq2 reveals gene expression of projection neurons. (**a**) UMAP plots of cortical excitatory neurons, colored by subclasses (*left*) and brains (*right*). (**b**)(**c**) The fraction of cells from each brain in each subclass (b) or cell type (c). (**d**) Overlap (dice coefficient) between all BARseq isocortical cell types and isocortical cell types in the ABC atlas (**e**) The locations of neuronal somata (black dots) that project to the indicated target cortical areas, from each subclass. Neurons that did not project to the target areas are shown in gray.

**Extended Data Figure 6.**
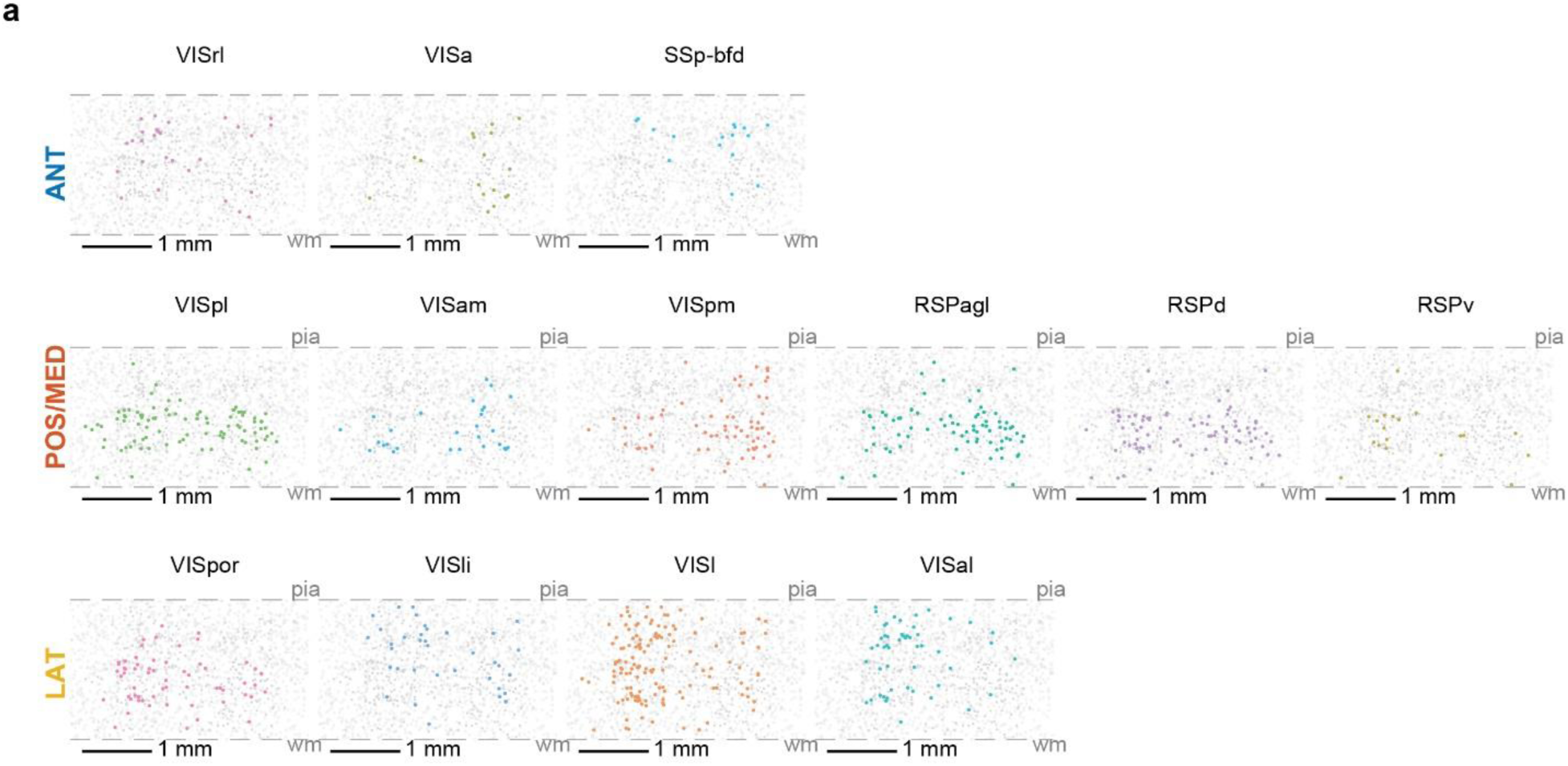
Laminar distribution of projection neurons is consistent with three pathways. “Coronal” flatmap views of neurons with the indicated projections. Each row includes areas for the same pathway. X axis indicates medial-lateral axis in flatmap coordinates, and y axis indicates depth.

## Methods

### Animals and surgery

Animal handling was conducted according to protocols approved by the Institutional Animal Care and Use Committee (IACUC) of the Allen Institute. Animals were housed in groups of 3–5 per cage on a 12/12 h light/dark cycle in an environmentally controlled room (40% humidity, 21 °C). Two C57BL/6J mice were used: one male (P64; animal 777552) and one female (P51; animal 824690).

Barcoded Sindbis virus libraries (CSHL MAPseq/BARseq core) were stereotaxically injected into VISp using coordinates derived from the Allen Reference Atlas. For animal 777552, virus was injected at five sites (AP, ML from Bregma, in mm): (−3.6, 2.5), (−3.6, 3.2), (−4.1, 2.2), (−4.1, 3.2), and (−3.1, 2.8). For animal 824690, the virus was injected at six sites: (−3.6, 2.5), (−3.6, 3.2), (−4.1, 2.2), (−4.1, 3.2), (−3.1, 2.8), and (−4.4, 3.0). At each site, 140 nL of virus was delivered at each of two depths, 0.7 and 0.3 mm below the brain surface; at the sixth site in animal 824690, the corresponding depths were 0.6 and 0.3 mm. Brains were collected approximately 20–24 h after viral injection.

### Tissue processing and sectioning

Approximately 24 h after Sindbis virus injection, mice were euthanized and transcardially perfused with ice-cold artificial cerebrospinal fluid (aCSF). Brains were removed, bi-sected along off the midline, and the injected hemisphere flash-frozen in a dry-ice-cooled isopentane bath and stored at −80 °C until sectioning. Brains were hemisected and cryosectioned coronally at 20 μm, starting from the posterior end and through VISp and the surrounding higher visual areas. Sections were collected into two interleaved series, such that consecutive sections were assigned alternately to the two series. One series was processed for in situ sequencing, corresponding to every other 20 μm section and therefore sampling one section per 40 μm of tissue (sequenced fraction = 0.5). A total of 112 hemi-coronal sections were processed for in situ sequencing across the two animals: 64 sections from animal 777552 and 48 sections from animal 824690.

### Axonal BARseq2 library preparation and in situ sequencing

BARseq libraries were prepared using a modified BARseq2 protocol (Sun et al., 2021), with the endogenous-gene panel and associated padlock probes previously developed and validated for cortex-wide cell-type profiling (Chen et al., 2025) (see **Supp. Table 1**). Briefly, slides were removed from −80 °C storage and immediately fixed in 4% paraformaldehyde (PFA) in PBS for 1 h at room temperature. Following a PBS wash, HybriWell-FL chambers were mounted over the tissue sections. Samples were dehydrated sequentially in 70%, 85%, and 100% ethanol for 5 min each, transferred to fresh 100% ethanol, and incubated for 1.5–3 h at 4 °C. Sections were then rehydrated by repeated washes in PBST (PBS containing 0.5% Tween-20).

Endogenous mRNAs were reverse-transcribed using 50 μM random N20 primer, while Sindbis barcode RNA was reverse-transcribed with 1 μM barcode-specific LNA primer as in BARseq2 (see **Supp. Table 1** for the sequence). Reverse transcription was performed in 1× RevertAid RT buffer containing 20 U μL⁻¹ RevertAid H Minus M-MuLV reverse transcriptase (Thermo Fisher Scientific), 500 μM dNTPs (Thermo Fisher Scientific), 0.2 μg μL⁻¹ BSA (New England Biolabs), and 1 U μL⁻¹ RiboLock RNase inhibitor (Thermo Fisher Scientific) at 37 °C overnight in a humidified chamber. The following day, samples were washed in PBST and the newly synthesized cDNA was covalently immobilized by incubation with BS(PEG)9 diluted 1:4 into PBST (for example, 200 μL BS(PEG)9 (Vector lab) stock plus 800 μL PBST) for 1 h at room temperature. Residual crosslinker was quenched by washing once with 1 M Tris-HCl, pH 8.0, followed by incubation in fresh 1 M Tris-HCl for 30 min. Samples were then washed twice with PBST. Endogenous transcripts were captured using the same 104-gene marker panel described previously (Chen et al., 2025). The panel contains up to 12 padlock probes targeting each gene; each padlock carries a 7-nt gene-identification index that encodes the identity of its target transcript. Gene padlocks were hybridized and ligated in 1× Ampligase buffer containing 100 nM of each padlock probe, 0.5 U μL⁻¹ Ampligase (Biosearch Technologies), 0.4 U μL⁻¹ RNase H (Qiagen), 1 U μL⁻¹ RiboLock RNase inhibitor, an additional 50 mM KCl, and 20% formamide. Samples were incubated for 30 min at 37 °C followed by 45 min at 45 °C. Without an intervening wash, Sindbis barcode cDNAs were captured by gap-filling padlock ligation as described for BARseq2. The reaction contained 100 nM barcode padlock probe, 50 μM dNTPs, 5% glycerol, 1 U μL⁻¹ RiboLock RNase inhibitor, 20% formamide, 50 mM KCl, 0.4 U μL⁻¹ RNase H, 0.001 U μL⁻¹ Phusion DNA polymerase (Thermo Fisher Scientific), and 0.5 U μL⁻¹ Ampligase in 1× Ampligase buffer. Samples were incubated for 5 min at 37 °C and 40 min at 45 °C. They were then washed twice with PBST and once with hybridization buffer (2× SSC, 10% formamide). An RCA primer was hybridized at 1 μM in hybridization buffer for 15 min at room temperature, followed by three 2-min washes in hybridization buffer and PBST washes.

Rolling-circle amplification was performed in two successive reactions. Samples were first incubated for 2 h at 40 °C with phi29XT polymerase. For 1mL mix, the phi29XT reaction contained 600 μL nuclease-free water, 200 μL 5× phi29XT reaction buffer, 100 μL 10 mM dNTP mix, and 100 μL phi29XT polymerase (New England Biolabs). The phi29XT mixture was then removed and replaced directly with conventional phi29 RCA mixture containing 1× phi29 buffer, 0.25 mM dNTPs, 125 μM aminoallyl-dUTP (Thermo Fisher Scientific), 0.2 μg μL⁻¹ BSA, 5% glycerol, and 1 U μL⁻¹ phi29 polymerase. RCA proceeded at room temperature overnight (>12 h). After RCA, samples were washed in PBST and the resulting rolonies were crosslinked with BS(PEG)9 in PBST for 1 h at room temperature. Excess crosslinker was quenched with 1 M Tris-HCl, pH 8.0, for 30 min, followed by PBST washes.

In situ sequencing was performed using Illumina Miseq nano v2 kits, following previously established protocol (Chen et al., 2025). Fluid exchange and image acquisition during sequencing were performed using a home-made automated fluidics and imaging system, built on a Nikon TI2E microscope with a Lumencor Celesta laser, a Prion NanoScan OP400 piezo objective scanner, Crest Xlight v3, and Photometrics Kinetix cameras. Filters and imaging conditions were the same as described previously (Chen et al., 2025).

### In situ sequencing data processing

In situ sequencing images were processed using a MATLAB pipeline as previously described (Chen et al., 2025). Briefly, maximum-intensity projections were denoised with Noise2Void(Krull et al., 2018), corrected for channel crosstalk and background, and registered across sequencing and hybridization cycles. Fields of view were stitched into whole-section images using MIST(Chalfoun et al., 2017), and gene rolonies were identified and base-called with BarDensr (Chen et al., 2021) against the padlock codebook. Somata were segmented with Cellpose(Stringer et al., 2021), masks were dilated by 3 pixels to accommodate residual registration error, and rolonies within each mask were assigned to the corresponding cell to generate the cell-by-gene count matrix. Duplicate cells in overlapping fields of view were removed by spatial matching within 10 µm. Sections were registered to the Allen Mouse Brain Common Coordinate Framework (CCF) using QuickNII and VisuAlign(Puchades et al., 2019) as previously described (Chen et al., 2025), providing CCF coordinates and anatomical annotations for each cell and rolony, as well as cortical streamline coordinates for cells in the isocortex.

Barcode images were registered to the gene-expression images through a shared reference channel and processed in the same whole-section coordinate system. Axonal barcode rolonies were detected after top-hat background subtraction (6-pixel radius), base-called in each sequencing cycle from the brightest of the four channels, and transformed into whole-section coordinates. For each rolony and cycle, base-call purity was defined as the intensity of the brightest channel divided by the Euclidean norm of the four channel intensities.

Soma barcodes were independently called from the registered barcode images using two approaches. First, channel intensities were averaged across all pixels within each cell mask and each cycle was assigned to the brightest channel. Second, calls were made using only high-purity pixels: pixels were retained when their third-lowest per-cycle purity exceeded 0.85, and cells containing more than 50 such pixels were called from their mean channel intensities. The high-purity-pixel call was used when available, with the whole-mask call used otherwise. Somata were classified as barcoded when the called sequence passed a sequence-complexity threshold and both the third-lowest per-cycle purity and third-lowest maximum-channel intensity exceeded 0.85 and 500, respectively. This automated calling was followed by manual annotation to remove cells that were mis-called as barcoded because of strong barcode labeling in neighboring cells and/or passing axons and/or dendrites. Using the third-lowest value allowed up to two sequencing cycles with poor signal without rejecting an otherwise high-quality barcode.

### Transcriptomically defined cell type clustering

Cells were typed as previously described (Chen et al., 2025). Cells with fewer than 20 total gene counts or fewer than 5 detected genes were excluded, and counts were normalized to counts per 10 and log2-transformed. Clustering was performed iteratively. At each round, we computed the first 30 principal components, constructed a shared-nearest-neighbor graph (k = 15; Jaccard edge weighting; approximate nearest neighbors), and partitioned the graph by Louvain community detection; UMAP (k = 15) was used for visualization. The first round separated glutamatergic, GABAergic, and non-neuronal cells; the second round was performed separately within the glutamatergic and GABAergic populations to identify subclasses; and the third round was performed independently within each glutamatergic subclass to identify clusters. At each round, clusters were annotated based on enrichment of curated marker genes, and clusters showing poor transcript detection rather than a distinct biological identity were excluded from cell-type analyses.

Animal 777552 was carried through all three rounds of clustering, whereas animal 824690 was clustered only at the first round. We then transferred subclass and cluster labels from 777552 to 824690 using hierarchical nearest-neighbor label transfer in a shared expression space. Both datasets were filtered and normalized identically, 50 principal components were fitted jointly across the two animals, and each 824690 cell was assigned the majority subclass among its 30 nearest 777552 neighbors. Cluster identity was then assigned by a second majority vote restricted to 777552 cells within the assigned subclass. Results were consistent across neighborhood sizes of 20, 30, and 50.

For some joint transcriptomic-projection analysis, we further filtered cells to requiring 30 reads/cell to ensure high-quality data.

### Barcode matching and projection-neuron quality control

Soma barcodes and axonal barcode rolonies were decoded independently and matched using Bowtie2, allowing at most one mismatch across the 15-nt barcode. Axonal rolonies were retained only if they matched a unique soma barcode, passed filters for over-represented library sequences and low sequence complexity, and belonged to a spatial cluster of at least 10 rolonies identified by DBSCAN. Rolonies within 500 µm of the parent soma were excluded to remove local axonal and dendritic signal.

To assess barcode-matching specificity, we calculated the minimum Hamming distance between each axonal barcode and the set of soma barcodes and compared the observed distribution with a null generated by independently shuffling each barcode position. For each neuron, we also calculated the number of distinct one-mismatch barcodes represented among its matched rolonies divided by the total number of matched rolonies. Neurons with a ratio ≥0.40 were excluded, because we expect that the majority of barcodes to be matched to one sequence.

The final projection-neuron cohort included neurons that (i) contained a manually verified soma barcode, (ii) had a soma located in VISp, (iii) contained at least 5 matched rolonies in annotated structures outside VISp, (iv) had these rolonies distributed across at least two sections, with no more than 80% on any single section, and (v) passed the 0.40 mismatch-ratio criterion. No soma barcode was shared between the two animals.

### Projection matrix and projection clustering

For most analyses, we counted matched ipsilateral rolonies in 18 target structures: 13 cortical targets (VISam, VISpm, VISrl, VISal, VISl, VISli, VISpor, VISa, VISpl, RSPd, RSPv, RSPagl, and SSp-bfd), dorsal lateral geniculate nucleus (LGd), lateral posterior nucleus (LP), lateral dorsal nucleus (LD), thalamic reticular nucleus (RT), and superior colliculus (SC). Cortical areas and LGd were assigned by their CCF parent structures, LP, LD, and RT by leaf structure, and SC as the union of its sensory and motor subdivisions. Somata with no rolonies in any of these targets were excluded from further analyses. Unless otherwise specified, projection profiles were normalized to sum to one across targets. For binary analyses, a neuron was considered to project to a target if at least 2 matched ipsilateral rolonies were detected in that target.

For projection clustering, row-normalized projection profiles were square-root transformed, such that Euclidean distance in the transformed space corresponds to Hellinger distance between the original projection distributions. Neurons were clustered by Ward hierarchical linkage for cluster numbers k = 4–20. For each k, we calculated the mean silhouette width, cluster-wise bootstrap Jaccard stability over 200 resamples using the clusterboot procedure, and adjusted Rand index relative to an alternative clustering based on cosine distance and average linkage. We selected the finest partition for which every cluster had a mean bootstrap Jaccard index ≥0.60 and contained at least 10 neurons, yielding k = 7. Clusters were subsequently ordered according to the target with the largest mean projection strength, with ties broken by the magnitude of that projection. For display, the stability analysis was additionally evaluated over k = 2–20. In neuron-by-target heatmaps, neurons were grouped by animals within each cluster to facilitate comparison between animals without affecting the clustering.

### Flatmap and side-view displays

Cortical soma locations were visualized using two standardized coordinate views. In flatmap views, cells were plotted according to their two tangential cortical-sheet coordinates. In side views, one tangential coordinate was plotted against normalized cortical depth, with the pia oriented upward, effectively unrolling the cortical sheet while integrating across the orthogonal tangential axis. Cells of interest were colored, and the remaining cells were shown as a subsampled grey anatomical reference. Axis limits, target colors, background subsampling, and masks for unannotated regions were held fixed across comparable panels.

Because the side view integrates across the anteroposterior extent of VISp, it was used to compare relative laminar ordering but not the absolute thickness of cell populations with measurements from individual physical sections.

### Comparison with bulk anterograde tracing

We compared the pooled BARseq projection profile with anterograde tracing data from the Allen Mouse Brain Connectivity Atlas. We retrieved ipsilateral projection_volume measurements from all wild-type VISp injection experiments (n = 15, experiment ID 100147853, 304586645, 307320960, 307557934, 309003780, 277712166, 277616630, 277714322, 307137980, 304585910, 309113907, 307593747, 100141219, 304564721, 304762965), summed projection signal over structures corresponding to each BARseq target and normalized each experiment across targets. To generate a bulk-equivalent BARseq profile without allowing neurons with high rolony counts to dominate, we averaged the row-normalized single-neuron projection profiles and renormalized the resulting mean profile.

Projection strengths were log10-transformed and compared using Pearson correlation. We quantified (i) the correlation between the pooled BARseq profile and the mean profile across the 15 atlas experiments and (ii) the correlation between each individual atlas experiment and the mean of the remaining 14 experiments, which provides an estimate of between-experiment reproducibility in the tracing dataset. Significance was assessed by permuting target identities 10,000 times.

### Comparison with single-neuron axonal reconstructions

#### Dataset and preprocessing

We compared BARseq projections with 169 fully reconstructed VISp neurons from a published fMOST single-neuron tracing dataset. Each reconstruction was separated into terminal arbor and fiber-of-passage components and resampled at approximately 50-µm intervals along the axon, so that sampled point density reflected axon length rather than reconstruction sampling density. For both datasets, analyses were restricted to ipsilateral axon outside VISp; the soma and local intra-VISp arbor were excluded.

#### Co-clustering

Each traced neuron was first converted into synthetic BARseq-like observations: for each synthetic observation, we drew a total rolony count, n, from the observed BARseq rolony-count distribution; we then sampled n points uniformly along the ipsilateral extra-VISp axon (terminal and passage equally likely per unit length) of a target neuron, then jittered by 100 µm to produce a BARseq-like observation. We drew the same number of synthetic observations as the number of BARseq neurons. The pooled matrix was clustered by Ward linkage on Hellinger distance across k = 2–30, as for the projection clustering. We report k = 26, the finest partition in which no cluster of ten or more neurons came from a single dataset, together with the joint projection matrix and the per-cluster composition. Because the subsampling is stochastic, the analysis was repeated over 25 realizations; a single-dataset cluster appeared in only one of the 25, so k = 26 is a conservative estimate of the consistency between the two datasets.

#### Match Consistency Score

To compare neurons between modalities at a spatial resolution finer than anatomical target areas, we defined a Match Consistency Score (MCS) between a query neuron and a reference reconstructed neuron. Spatial correspondence was evaluated with the kernel

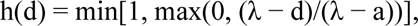

where d is Euclidean distance in CCF space, a = 100 µm, and λ = 1,000 µm. Query coverage was defined as the mean, over query observations, of the maximum match to either terminal arbor or fiber of passage, with fiber-of-passage matches down-weighted by ρ = 0.5. Reference coverage was defined as the axon-length-weighted mean match between the reference terminal arbor and its nearest query observation. Fiber of passage was excluded from reference coverage to avoid penalizing long-range neurons for the spatial extent of a single traversing axon. Because reference coverage depends on the number of observations in the query neuron, it was normalized to a rolony-count-matched ceiling estimated from 100 Monte Carlo draws at each count. MCS was defined as the harmonic mean of query coverage and reference coverage.

For every BARseq and reconstructed neuron, we identified the highest-scoring reference reconstruction, excluding self-matches. As a spatial negative control, BARseq neurons were displaced by 1.5 mm and rematched against the same reference library. Robustness was evaluated by independently varying λ, ρ, coordinate jitter, and whole-target dropout.

#### Soma-depth prediction

As an independent test of whether axonal similarity contains information about soma position, we predicted cortical depth using the top three reconstruction matches for each neuron. Only reference matches with MCS ≥0.80 were retained, and their soma depths were averaged; neurons without qualifying matches received no prediction. BARseq neurons with soma depth <40 streamline units were excluded from this analysis because few traced neurons were in the same depth range. The same readout was applied to a control: leave-one-out matching among traced single neurons, which removes the effect of sparse BARseq sampling.

### Detection sensitivity and cross-platform comparison

#### Detection sensitivity

We estimated axonal detection sensitivity by comparing branched axon length in reconstructed neurons with barcode rolony counts for the same projection. We used the thalamic collateral of superior-colliculus-projecting neurons as the reference projection because it was robustly observed in the tracing dataset (73 of 76 SC-projecting neurons), and was a weaker projection that did not saturate detection at the sensitivity of axonal BARseq. The ratio of median branched axon length in the tracing dataset to median thalamic rolony count in BARseq provided an estimate of micrometers of axon per detected rolony. Because every other 20-µm section was sequenced, the corresponding physical detection limit was taken as one-half of this value.

We next simulated rolony detection from reconstructed axons over sensitivities spanning 220 (the measured sensitivity after accounting for sampling frequency) – 814 µm (detection floor based on the TH collaterals of SC-projecting neurons) of axon per rolony. Traced neurons were assigned to broad transcriptomic classes from their predicted MET identities in the order ET, CT, L6b, NP, Car3, and IT/RSP. For each class, we measured the fraction of neurons for which the expected signature projection would be detected: extra-VISp isocortical terminal arbor for IT neurons, SC for ET neurons, and thalamus for CT neurons. Only terminal arbor was included.

#### Cross-platform comparison

We compared sensitivity across barcoded-connectomics platforms using detection of thalamic collaterals in SC-projecting neurons as a common benchmark. The comparison included the present dataset, the original axonal BARseq dataset, a bulk MAPseq/BARseq dataset in which projection sites were sequenced, and an AAV-based Synapse-seq dataset.

For the primary comparison, an SC-projecting neuron was required to contain at least 5 detected units in SC, where a unit was a matched rolony for in situ datasets, a molecule count for MAPseq, and a distinct viral tag for Synapse-seq. Because this criterion yielded only 1 L5 ET cell in the Synapse-seq dataset, a ≥1-unit threshold was used instead. Each dataset was paired with an internal negative-control population containing zero SC units: neurons defined by contralateral cortical projections for the two auditory-cortex datasets, neurons with superficial somata for the present dataset, and glia for Synapse-seq. Proportions were reported with Wilson confidence intervals, pairwise proportions were compared by Fisher’s exact test with Benjamini–Hochberg correction, and the distributions of thalamic counts among SC-positive neurons were examined directly.

### Retinotopy: reference and projection-inferred maps

#### Functional retinotopy reference

Because retinotopic organization varies across animals, we constructed an average reference map from published intrinsic-signal imaging (ISI) data from the Allen mouse brain connectivity atlas. We used brains that could be accurately registered to CCF and excluded brains with satellite patches in higher-visual-area fits, resulting in 283 specimens. For each specimen, azimuth and altitude phase maps were transformed into the cortical flatmap, and the median value per pixel was used to construct the mean maps. Visual field sign was calculated from the phase gradients in each animal’s native acquisition coordinates before warping and then averaged across animals. Because the ISI data were not calibrated to visual angles, they were calibrated to visual angles by comparing to functional retinotopy map from (Waters et al., 2019): we registered the mean map to the field sign map in flatmap space, then regressed the azimuth and altitude values from the Waters et al. data to the mean maps. This map was used to assign retinotopic coordinates to neurons. To compare to projection-defined maps, the reference brains were smoothed with a Gaussian kernel with σ = 20 pixels for azimuth and altitude, and σ = 6 pixels for field sign, then combined to build smoothed reference maps.

#### Projection-inferred retinotopic maps

Projection-inferred maps were constructed by calculating, at each cortical flatmap location, the rolony-weighted mean azimuth or altitude of the VISp somata giving rise to nearby axonal rolonies. Flatmap pixels were included only when they exceeded a minimum rolony-count threshold (100 rolonies within a 40 pixel radius in flatmap space), and inferred values were rescaled to the range of the corresponding functional reference. Visual field sign was calculated from the inferred azimuth and altitude maps using the same phase-gradient procedure as for the ISI data.

Projection and functional maps were compared over a common spatial mask restricted to the target areas, pixels with valid ISI measurements in all contributing brains, and pixels defined on the projection side. Agreement was quantified by pixel-wise Pearson correlation separately for azimuth and altitude. As a reference for biological variability, each ISI brain was correlated with the average of the remaining brains in a leave-one-out analysis.

#### Retinotopic and positional organization of projections

For visualization of corticothalamic topography, the two corticothalamic projection clusters from the k = 7 partition were plotted in the cortical side view. Their thalamic rolonies were then plotted in coronal CCF coordinates and colored by the calibrated visual-field position of the parent soma, allowing the organization of VISp inputs within LGd and LP/LD to be related directly to source retinotopy.

To assess positional biases in cortical target selection, neurons were plotted on the VISp flatmap according to whether they projected to each target, using a threshold of at least 2 ipsilateral rolonies. Analyses were performed for the full cohort of neurons and separately within each of the four IT subclasses.

For each target, we tested whether projecting and non-projecting neurons differed along the mediolateral or anteroposterior axis of VISp. VISp was divided at the median along each axis, and association between cortical position and projection status was tested using a Cochran–Mantel–Haenszel test stratified by animal. P values were corrected across targets and axes using Benjamini–Hochberg FDR.

### Co-projection structure and models of target grouping

#### Co-projection enrichment and retinotopy control

We first tested the 50 bifurcation, trifurcation, and quadfurcation motifs previously reported across six higher visual areas. For each motif, the observed number of neurons carrying the corresponding projection footprint was compared with the expectation derived from the marginal projection probabilities. P values were corrected across motifs using Holm–Bonferroni correction.

We next quantified pairwise co-projection enrichment as the observed probability of projecting to targets i and j relative to the expectation P(i)P(j). To determine how much pairwise enrichment could be explained by soma position in VISp, we constructed a retinotopy-conditional null. For each neuron and target, projection probability was estimated from soma position using a leave-one-out Nadaraya–Watson kernel regression fitted separately within each animal, with a cross-validated bandwidth of 300 µm. The expected co-projection probability was then calculated as the mean across neurons of the product of the two target-specific probabilities. Residual enrichment relative to this expectation represents co-projection structure not explained by soma position.

#### Models of target grouping

We modelled binary projection footprints using pairwise maximum-entropy (auto-logistic) models. The log-odds of projection to each target contained a target-specific intercept, coupling terms describing its relationship to the other targets projected to by the same neuron, and a fixed per-neuron offset given by the retinotopy model above, so that any coupling structure must improve on retinotopy rather than recapitulate it. The partition function was evaluated exactly by enumerating all possible footprints, giving exact maximum-likelihood fits; models were fitted by Newton’s method with a ridge penalty on the coupling terms only, removed for held-out scoring.

We compared model classes at matched complexity. In the ring model, pairwise coupling was a constant plus a term proportional to the cosine of the angular separation between targets around VISp on the cortical flatmap; a variant using unwrapped angular distance was also fitted, and the better-fitting of the two served as the reference against which all other models are reported. In the block model, pairwise coupling took one value for targets within the same group and another for targets in different groups. Both classes therefore contain exactly two coupling parameters, at any number of groups, so their likelihoods compare directly with no complexity correction. For reference, we also fitted a fully unconstrained pairwise model and a model containing no pairwise coupling. Target angles were defined from CCF flatmap centroids relative to the VISp centroid, with the angular branch cut placed in the largest gap between targets; the cut affects only the unwrapped variant, as the cosine form is invariant to it. Block assignments were searched exhaustively under the constraint that each group form a spatially connected set on the flatmap, using shared areal borders together with a closure edge joining the two targets at the extremes of the angular ordering. Searching two to four groups yielded 80 candidate bi-partitions, 620 tri-partitions and 1,895 quad-partitions, 2,595 candidates in total. Significance was calibrated using a max-statistic parametric bootstrap: two hundred datasets were simulated and the complete partition search was repeated on each, generating the null distribution of the best-of-scan statistic within each group-number family and across all of them.

### Mapping transcriptomically defined cell types to the Allen Brain Cell atlas

Cortical excitatory cells were mapped to the Allen Brain Cell (ABC) whole-mouse-brain transcriptomic taxonomy using k-nearest-neighbor majority voting (k = 15) over genes shared between the BARseq panel and reference dataset. *Slc17a7* and *Gad1* were excluded from the matching features. To account for systematic gene-specific differences in detection between platforms, each reference gene was multiplied by the ratio of its mean expression in the BARseq dataset to its mean expression in the reference dataset. Expression profiles were then z-scored within each cell before nearest-neighbor matching. Correspondence between BARseq and reference labels was summarized by a Dice-overlap matrix. Reference labels with no overlap exceeding a Jaccard index of 0.05 were omitted from visualization, and three reference clusters outside the cortical taxonomy were excluded. Mapping was performed both across the full isocortical excitatory taxonomy and after restricting both query and reference populations to visual cortex. As a sensitivity analysis, the mapping was repeated after Harmony integration in principal-component space; the per-gene rescaling approach was retained for the reported results because it performed better at the isocortical scope. For L4/5 IT neurons, mappings were additionally summarized separately for each of the eight L4/5 IT clusters.

### Correspondence between transcriptomic type and projection

For each transcriptomic type, we calculated the mean row-normalized projection profile across the 18 targets. Analyses were restricted to neurons with at least 30 detected gene counts and to transcriptomic types represented by more than 10 barcoded neurons. We also calculated the transcriptomic composition of each projection cluster and, conversely, the projection-cluster composition of each transcriptomic subclass.

To validate the correspondence between cell types and projections, we examined the spatial distribution of the corresponding cell types in the ABC MERFISH dataset, based on cell type correspondence established by the knn mapping described above.

### Position versus gene expression in L4/5 IT neurons

#### Variance explained by spatial and transcriptomic features

For each of the 13 cortical targets, we fitted cross-validated models predicting the binary projection call from (i) tangential soma position on the cortical flatmap, (ii) the first 15 principal components of gene expression, or (iii) cortical depth. Identical neurons, cross-validation folds, and random seeds were used for all three predictors. We report the cross-validated R² for each model and the difference between the tangential-position model and each alternative. Targets were additionally classified as proximal or distal to VISp and the distributions of model performance compared by Wilcoxon rank-sum test.

#### Position-matched null for expression analyses

Because the canonical projection groups differ in soma position, analyses relating projection group to gene expression within a subclass were evaluated against a position-matched null. For each labeled neuron, surrogate labels were sampled from the set comprising that neuron and its four nearest neighbors in tangential flatmap coordinates within the same animal, without replacement across labeled neurons, thereby preserving the spatial distribution of the projection groups; the analysis was repeated with eight and twelve neighbors as a sensitivity check.

Against this null, we ran a scan over partitions of the transcriptomic-cluster ladder ordered by median expression pseudotime, using the minimum Fisher’s exact-test p value as the statistic and repeating the complete cut search for each null replicate, so that the resulting p is corrected for cut selection as well as for position; the equivalent scan over a depth-ordered ladder was computed in the same family. Comparisons with insufficient spatial overlap between projection groups were classified as inseparable rather than assigned a p value.

For expression-axis analyses, principal components were fitted using the entire VISp L4/5 IT population and barcoded neurons were projected into this basis. Projection groups were treated as non-exclusive, so neurons projecting to multiple groups contributed to each relevant group. Separation was quantified as the AUROC along PC1 and PC2 and as the cross-validated AUROC of a ridge-logistic classifier fitted to the first two components, with five-fold cross-validation; AUROCs are reported oriented so that values above 0.5 indicate higher scores in the posteromedial group. Significance was assessed by permuting projection-group labels within animal (5,000 permutations), and reported values are uncorrected.

### Statistics

Statistical tests are specified with each analysis. Group comparisons used Wilcoxon rank-sum or signed-rank tests, Kolmogorov–Smirnov tests, Fisher’s exact tests, or Cochran–Mantel–Haenszel tests stratified by animal, as appropriate; associations were assessed by Pearson or Spearman correlation. Classifier and one-dimensional separation performance were quantified by area under the receiver-operating-characteristic curve. Model performances were cross-validated, except for the comparison of collateral projection models, which is reported in sample. In this analysis, significance was assessed against a parametric bootstrap in which every model was refitted on each simulated dataset, so that any advantage conferred by fitting is present in the null as well as in the statistic.

Where a model, partition or cut point was selected from a search, significance was assessed by a max-statistic permutation or parametric-bootstrap procedure in which the complete selection process was repeated for every null replicate. Model classes were compared at matched numbers of free parameters, so their likelihoods are directly comparable without a complexity penalty. Other permutation tests were stratified by animal unless stated otherwise. Permutation and bootstrap p values were computed as (1 + the number of null replicates reaching the observed statistic)/(1 + the number of replicates), so the smallest attainable value is set by the number of replicates. Multiple comparisons were controlled by Holm-Bonferroni or Benjamini-Hochberg as indicated. Proportions are reported with Wilson confidence intervals.

## Data availability

Processed axonal BARseq2 dataset is deposited at Synapse.org (https://doi.org/10.7303/syn77214713).

## Code availability

Analysis code is provided at Synapse.org (https://doi.org/10.7303/syn77214713). BARseq data processing pipeline is available at github (https://github.com/chenxy877/barseq-processing.git).

## Supplementary materials

*Supplementary Table 1: List of genes in the gene panel and oligos used for BARseq*

## Acknowledgements

We thank Li Yuan and Fangming Xie for insightful discussions; and Gregory Horwitz, Angela Fan, James Bourne, Hongkui Zeng for feedback on the manuscript. This work was supported by the National Institutes of Health (NIH) [U01NS132161, R01MH133181, DP2MH132940 to X.C.]. The authors thank the Allen Institute founder, Paul G. Allen, for his vision, encouragement, and support. The findings and conclusions presented in this paper are those of the author(s) and do not necessarily reflect the views of the NIH or the

U.S. Department of Health and Human Services.

## Author contributions

Conceptualization: X.C.; Investigation: M.M., S.C., S.K., A.A., O.H., K.N., B.O., A.O., D.R., J.W., X.C.; Methodology: M.M., M.C.P.R., A.Z., J.W., Y.I., X.C.; Resources: M.M., M.C.P.R., X.C.; Validation: M.M., M.C.P.R., A.Z., X.C.; Data curation: M.M., S.C., X.C.; Software: M.M., M.C.P.R., A.Z., X.C.; Formal analysis: M.M., X.C.; Visualization: M.M., M.C.P.R., X.C.; Writing - original draft: M.M., X.C.; Writing - review and editing: M.M., X.C.; Supervision: J.A., M.R., A.W., Y.I., X.C.; Project administration: X.C.; Funding acquisition: X.C.

